# Whole-genome sequencing across space and time reveals impact of population decline and reduced gene flow in Florida Scrub-Jays

**DOI:** 10.1101/2024.11.05.622154

**Authors:** Tram N. Nguyen, Elissa J. Cosgrove, Nancy Chen, Natasha Lehr, Mitchell G. Lokey, Felix E. G. Beaudry, Sarah W. Fitzpatrick, Reed Bowman, Karl E. Miller, John W. Fitzpatrick, Andrew G. Clark

## Abstract

Whole-genome sequence data is proving to be highly informative about the past demography of free-living populations, and in the context of endangered species, it can provide a quantification of the genetic risk posed by reduced genetic diversity and inbreeding. Prior to 1920, the Florida scrub-jay (*Aphelocoma coerulescens*) had been widespread across Florida, but with the expansion of agriculture and human habitation, its population has declined by 95%, resulting in fragmentation into semi-isolated subpopulations. By sequencing 241 individuals sampled from five different regions and across two time points, this study quantifies a greater magnitude of loss of genetic diversity and greater levels of inbreeding in smaller and more isolated subpopulations. Consistent with population genetics theory, reduction in population size results in a dramatic loss of rare alleles, skewing the site frequency spectrum far from the expected equilibrium. Increased inbreeding in the smaller, more remote subpopulations is especially evident in the increased size and number of runs of homozygosity. The Florida scrub-jay displays limited dispersal, and habitat fragmentation has greatly reduced the magnitude of gene flow in the past 30 years, resulting in further decline of genetic diversity, especially in the peripheral populations. Analysis of these data is informative in guiding conservation efforts to retain genetic diversity and minimize the consequences of inbreeding in the Florida scrub-jay.

**Highlights:**

- Five regional populations show distinct degrees of population isolation and decline.
- There has been commensurate loss of genetic diversity, skewed site frequency spectra, reduced migration, and increased inbreeding (*F_ROH_*).
- As many state-wide populations decline, the smaller, more remote populations provide a glimpse into the future and a testbed for remediation approaches.

## Introduction

Characterizing spatial and temporal patterns of genetic diversity and gene flow among populations undergoing decline is essential for predicting the future of vulnerable species, and for exploring fundamental questions about contemporary evolution. Extrinsic pressures that reduce population size, such as habitat loss, provide opportunities to test our understanding of basic population genetic principles as they apply to free-living populations. Small, isolated populations are expected to lose genetic diversity owing to genetic drift and reduction of gene flow. Loss of standing genetic variation, in turn, can reduce the adaptive potential of species facing environmental perturbations and reduce their resistance to disease (Anderson & May, 1986; Pearman & Garner, 2005; Spielman et al., 2004; Tarpy, 2003; Whiteman et al., 2006). Genetic drift also can elevate the frequency of deleterious alleles, magnifying harmful consequences of inbreeding and potentially driving small, declining populations to extinction (Frankham, 2005; Frankham et al., 1999; Lande, 1994; Lynch et al., 1995a, 1995b; Gilpin & Soule, 1986; Mills & Smouse, 1994). Thus, recording the genetic diversity and degree of inbreeding within populations undergoing varying levels of decline may prove invaluable not only for understanding the consequences of population declines but also how quickly genomic degradation can occur through the accumulation of deleterious variants.

Current levels of genetic variability are shaped both by recent population trajectories and by deeper demographic history, especially as past population bottlenecks can influence the ability of populations to purge deleterious variants, thereby affecting population persistence (Grossen et al., 2020; Robinson et al., 2016). Inferring the timing and scale of past bottlenecks thus provides essential context for understanding the genomic consequences of population decline in endangered species. Furthermore, characterizing the demographic past of populations lays the foundation for exploring evolutionary questions about selection in the future (Charlesworth, 1994, 2009; Kimura, 1968). By examining populations with varying demographic histories and degrees of isolation, we can evaluate the impact of these fundamental processes on genetic variability and inbreeding. Altogether, attributes such as genetic diversity, population structure, gene flow, and demographic histories are critical for informing immediate and long-term conservation efforts, including genetic rescue (Kyriazis et al., 2021).

Neutral molecular markers such as microsatellites and ddRAD have informed numerous conservation genetic studies in wild species (Wilder et al., 2020; Hardy et al. 2021; Modi et al. 2021; Gu et al. 2021), but recent advances in whole-genome sequencing (WGS) have unlocked vastly greater capacity to study genomic variation, revealing remarkably detailed insights into the genomic consequences of population decline and inbreeding (reviewed in Hohenlohe et al., 2021; Allendorf 2017). Notably, WGS best resolves genomic segments that are identical-by-descent (IBD) across individuals, and also accurately detects runs of homozygosity (ROH). Individuals share IBD segments via inheritance of homologous chromosome segments from a common ancestor, and inbred individuals may manifest these IBD chromosomal regions as contiguous tracts of homozygosity in DNA sequences (ROH). IBD segments provide information about relatedness and gene flow, while ROH provide useful metrics of inbreeding (Leitwein et al., 2020). Furthermore, because both IBD segments and ROH are fragmented over time by recombination (Franklin, 1977), they can be used for inferring time since common ancestry and timing of inbreeding events. Finally, haplotype-based models using IBD segments allow for robust inference of very recent demographic histories (*i.e.*, within the last few generations), and can harness information about linkage disequilibrium (LD) and recombination rates for accurate estimates of effective population sizes (Delaneau et al., 2019; Harris & Nielsen, 2013; Lawson et al., 2012; Marchi et al., 2021; Ralph & Coop, 2013; Terhorst et al., 2017).

The Florida Scrub-Jay (*Aphelocoma coerulescens*; hereafter FSJ) provides an ideal and timely system for exploring spatiotemporal patterns of genomic diversity in a species facing widespread habitat loss and population decline. FSJ is a federally Threatened, cooperatively breeding, nonmigratory bird species endemic to patches of fire-maintained oak scrub unique to the Florida peninsula (Woolfenden and Fitzpatrick, 1984). Genetically distinct metapopulations, shaped by a complex paleoclimatic history, including periodic expansions and bottlenecks resulting from glacially-related wet-dry cycles (Coulon et al., 2008; Hine et al. 2017), have undergone severe population declines and local extirpations resulting from extensive anthropogenic habitat conversion in recent history (Boughton & Bowman, 2011; Woolfenden & Fitzpatrick, 2024). In particular, large-scale commercial agriculture (especially citrus) and human colonization since the late 1800s dramatically accelerated habitat fragmentation and loss. The species today is estimated to be below 10% of its pre-1900 size (Boughton & Bowman, 2011). Variously sized FSJ populations persist, mainly where their unique habitat is protected and actively managed, with emphasis on prescribed burning (Breininger et al., 2018; Breininger & Oddy, 2004; Fitzpatrick & Bowman, 2016). These populations, several of which have been intensively studied for decades, offer an extraordinary opportunity to contrast the genomics of conspecific populations having substantially different sizes, demographic histories, degrees of isolation, and population trajectories.

A range-wide evaluation of FSJ population structure using microsatellites based on over 1,000 samples collected between 1992 and 2003 from found the landscape could be best characterized by 21 metapopulations (Coulon et al. 2008). In 2018 (4-5 generations after the first samples), we collected modern samples from five of these populations (Figure 1), carefully chosen to represent a range of sizes and trajectories. These focal populations are as follows, from largest to smallest:

1. Ocala National Forest (ONF; Figure 1B), in north-central Florida, hosts the largest remaining FSJ population (Miller and Shea 2019, USFWS 2019), with at least 1,000 family groups (∼3,000 individuals).
2. Archbold Biological Station (ABS) is a private, well-managed ecological preserve in south-central Florida (Turner et al., 2006; Weekley et al., 2008) which harbors more than 200 FSJ family groups (∼600+ individuals). This population has been intensively studied since 1969 (Fitzpatrick and Bowman, 2016), yielding a 16-generation pedigree, detailed ascertainment of immigration rates (Chen et al., 2016), and a linkage map (Romero et al., 2024).
3. Placid Lakes Estates (PLE), located 10 km northwest of ABS within the same genetic metapopulation (Coulon et al., 2008), is a densely populated residential subdivision. FSJs numbered in the hundreds here as recently as 1990 but are now reduced to a small handful of family groups (Nguyen et al., 2022).
4. Jonathan Dickinson State Park (JDSP), on the southeastern coast, has more than 1,200 ha of suitable FSJ habitat, capable of supporting 70-100 FSJ family groups. However, as of 2013, it had fewer than 30 (90 individuals). Without neighboring populations, JDSP is vulnerable to inbreeding depression and is a priority site for genetic rescue efforts.
5. Savannas Preserve State Park (SPSP), north of JDSP, is one of Florida’s smallest protected populations (∼20-30 individuals) and occasionally receives immigrants from other tiny populations located farther north.

**Figure 1.**
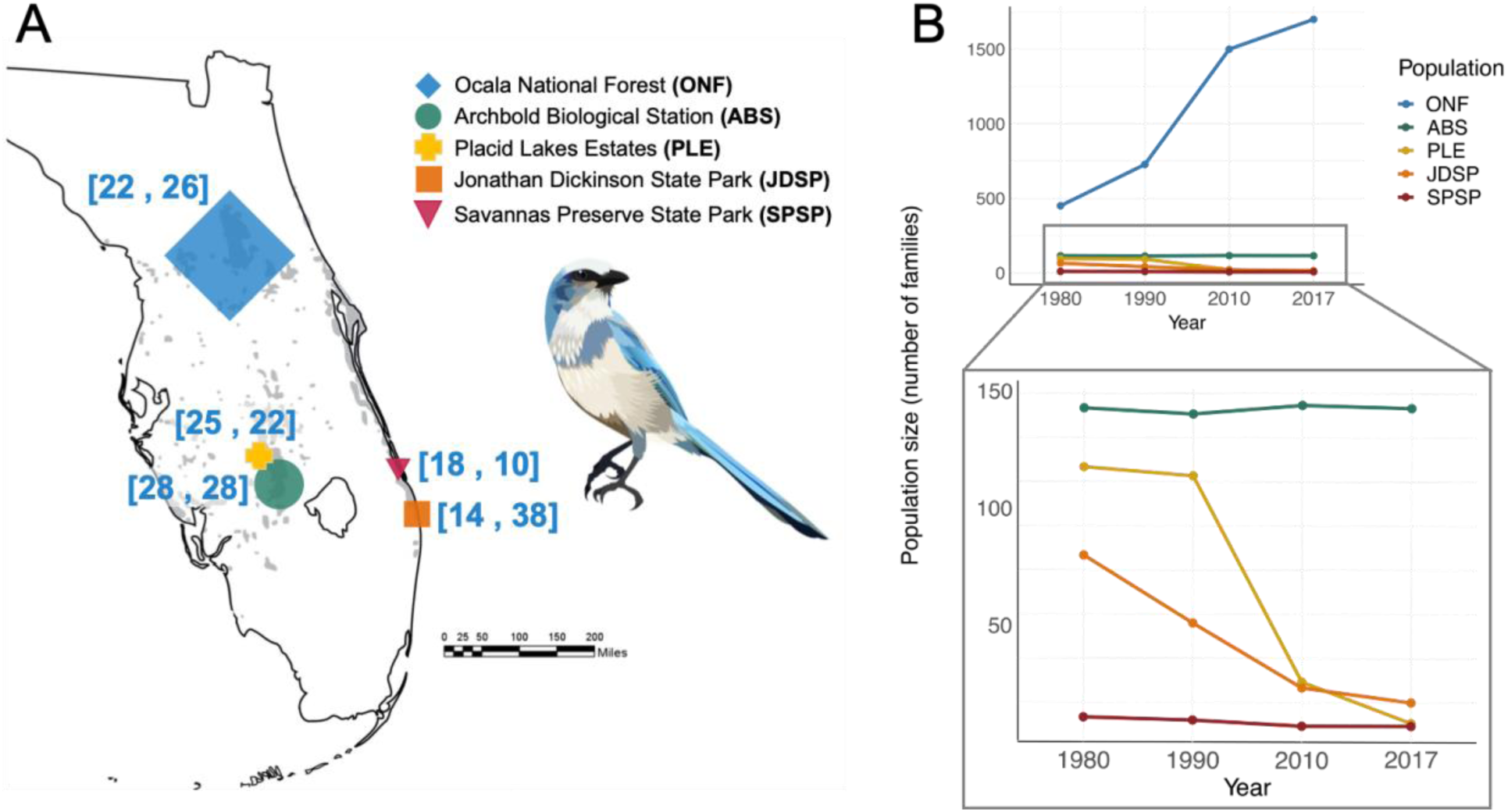
**A)** Florida Scrub-Jay range distribution and sample map. Gray areas represent the current species’ distribution and symbol shapes represent focal populations in the present study. Symbol sizes correspond to relative current census sizes. The first and second number in each bracket represent historic and contemporary sample sizes for this study, respectively. **B)** Estimated census sizes of focal populations using anecdotal and census observations emphasizing the large, stable nature of ONF and ABS, the sharp, recent declines of PLE and JDSP, and the persistently small population of SPSP.

Our multi-decadal, whole-genome sequences of 241 individuals provide an exceptional resource to comprehensively evaluate the consequences of population decline on genetic variability, inbreeding, and relatedness across spatial and temporal scales. Moreover, combining both SFS- and IBD segment-based approaches enables a detailed reconstruction of population dynamics across deeper and recent timescales. Our research serves as a critical evaluation of a precipitously declining species that is immediately informative for conservation management, and establishes invaluable genomic resources and groundwork for future studies investigating fundamental evolutionary and population genetics processes on contemporary timescales.

## Results

### Population structure and genetic ancestry

The results of Principal Components Analysis show clear partitioning of the geographically separate populations (Figure 2A), consistent with the genetic population structure previously inferred from microsatellite data (Coulon et al. 2008). The neighboring coastal populations of JDSP and SPSP show a degree of genetic divergence despite their limited geographic separation (∼30 km), likely due to unsuitable intervening habitat (e.g., dense residential areas) (Stith et al. 1996). Samples from ABS and PLE appear as a single genetic group in PCA space, reflecting close genetic similarity and modest levels of gene flow between the subpopulations, which is also supported by banding data (Nguyen et al. 2022).

**Figure 2.**
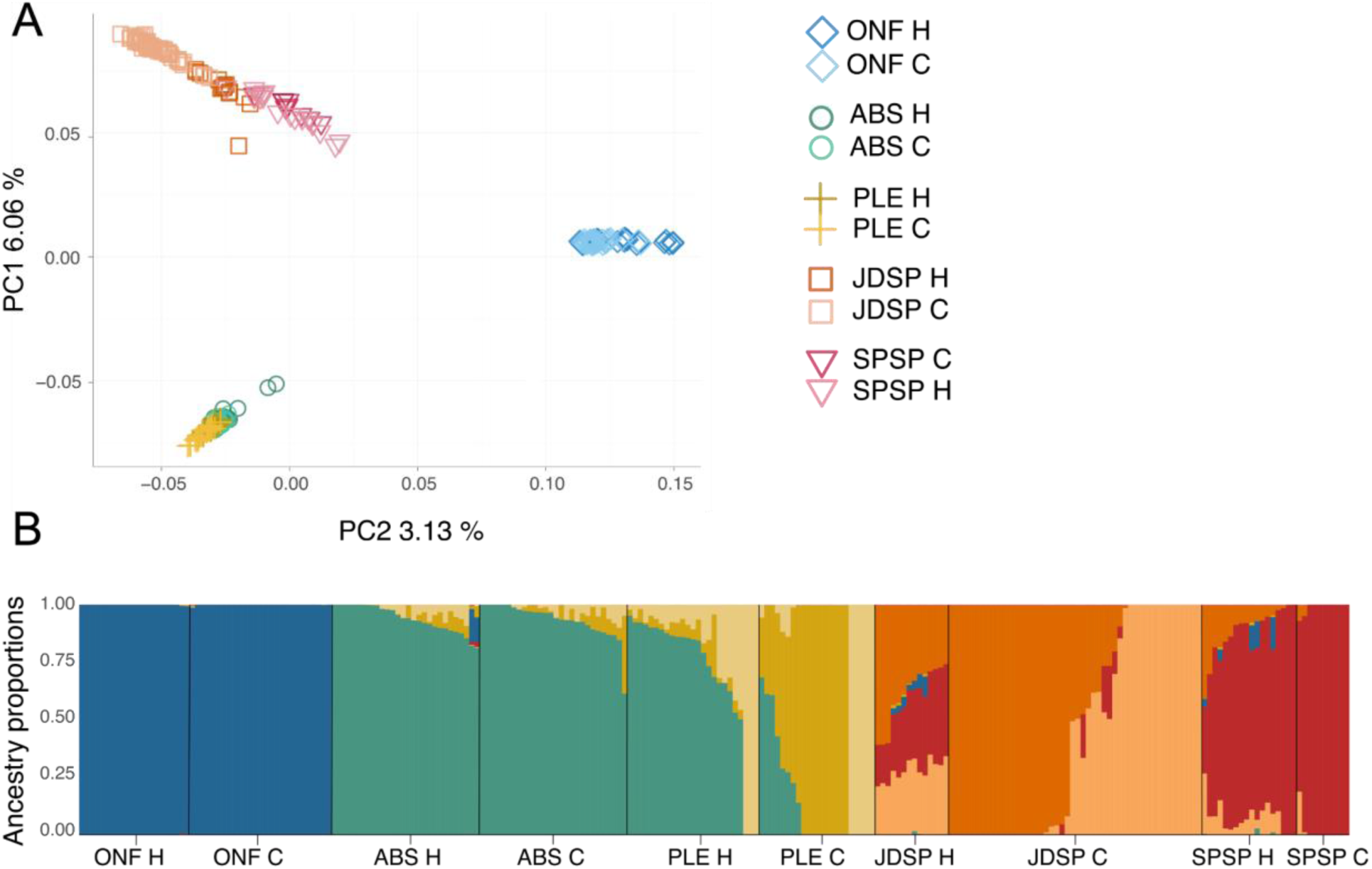
**A)** Principal component analysis (PCA) using 8,901,142 autosomal, biallelic SNPs of high-confidence and quality. Each point represents a breeding individual included in the main study (*n*=241). **B)** Composition of genetic ancestry of the 241 individuals. Each individual corresponds to a column, with the proportion of each color reflecting the proportion of ancestry assigned to a genetic group. The optimal number of genetic clusters (*K*=7) was identified by minimum cross-validation error. For example, the genomes of individuals in ONF are almost completely derived from an ONF-specific lineage (colored blue), as opposed to PLE individuals, which possess a pervasive admixture of ABS ancestry. The “H” and “C” following each population label in the legend correspond to the historic and contemporary sampling groups for each population, respectively.

To visualize fine-scale genetic structure between samples across populations and time, we plotted the ancestral composition of our 241 breeders using the optimal cluster number, *K*=7 (Figure 2B). Our results were consistent with our PCA clustering and known metapopulation structure. ONF samples showed distinct ancestry, *i.e.* very little ancestry shared with any of the other four populations across both sampling periods (average shared ancestry with other populations within both ONF groups = 1.94 × 10^−3^).

ABS samples also had most of their ancestry assigned to a distinct ABS lineage, however, nearly all individuals in ABS also shared a small proportion of their ancestry with PLE (5.00 × 10^−2^ in historic and 6.70 × 10^−2^ in contemporary). This supports evidence for historic and ongoing gene flow between these populations, with more PLE individuals moving into ABS in contemporary times as suitable habitat in the suburban PLE declined. In historic times, PLE birds harbored extensive ABS ancestry (mean = 0.70), but the proportion of ancestry from ABS was greatly decreased in PLE contemporary samples (mean = 0.14), consistent with a reduction in gene flow from ABS to PLE. Curiously, two distinct genetic lineages are present in the PLE contemporary samples. This could occur when populations become more fragmented and start to differentiate from a once-panmictic population, or from more recent immigration. The same is seen in JDSP contemporary samples, where two different lineages also exist, likely indicative of genetic substructure within this population caused by isolation-by-distance (Aguillon et al., 2017) during historic times when large populations existed both north and south of JDSP. Indeed, greater SPSP co-ancestry was found in historic JDSP samples (mean = 0.30) than in JDSP contemporary samples (0.016). Finally, SPSP ancestry is relatively distinct in its population, with more JDSP co-ancestry in historic SPSP samples, further suggesting more admixture between these populations in the past. Notably, some ONF and ABS ancestry is also revealed in the historic east coast populations, indicative of deeper historic gene flow between large, stable populations and long-distance sink populations. The overall picture reveals gene flow among populations of Florida scrub-jays has considerably declined during recent FSJ generations.

### Genetic diversity

We compared nucleotide diversity (π) and the genome-wide heterozygosity per individual *(H_0_)* across sampling periods and among populations (Table 1; Table S1). Nucleotide diversity, a commonly used metric for genetic diversity, is calculated as the average number of nucleotide differences per site between all possible pairs of DNA sequences in a population sample. The genome-wide heterozygosity (*H_0_*) is the proportion of all nucleotides that are heterozygous, calculated for each individual. These two metrics can differ if the population deviates from Hardy-Weinberg proportions, and in our sample, they are generally very close. Although not statistically significant, *H_0_* increased in ONF and ABS across the sampling interval, but dropped in the three small, declining populations at PLE, JDSP, and SPSP (Figure 3A; Table 1; Table S2). Interestingly, ABS Historic showed the lowest levels of *H_0_* and π, although this was not significantly different from PLE or JDSP Historic. In contemporary times, ONF had the highest level of *H_0_* and π among populations, followed by ABS. PLE Contemporary dropped to the lowest level of *H_0_* and π, and is now significantly different from ABS Contemporary despite being in the same metapopulation (Table S3).

**Table 1.** Summary statistics across populations. Values represent the means and standard errors of each statistic, if relevant.

| Population group | Sample size (n) | Mean genomewide heterozygosity ($H_o \times 10^{-3}$) | Mean nucleotide diversity ($\pi \times 10^{-3}$) | Mean Tajima's D | Mean inbreeding (FROH) | Mean number of ROH | Mean ROH Length (kb) | Proportion long ROH (> 5 Mb $\times 10^{-4}$) | Mean relatedness (pairwise IBD) |
| --- | --- | --- | --- | --- | --- | --- | --- | --- | --- |
| ONF Historic | 22 | 1.86 ± 0.04 | 1.72 | 0.578 | 0.106 ± 0.013 | 99.75 ± 11.43 | 47.54 ± 156.24 | 1.24 | 0.045 ± 0.035 |
| ONF Contemporary | 26 | 1.87 ± 0.04 | 1.78 | 0.439 | 0.109 ± 0.012 | 93.91 ± 4.04 | 43.96 ± 139.74 | 1.10 | 0.010 ± 0.009 |
| ABS Historic | 28 | 1.66 ± 0.08 | 1.54 | 0.868 | 0.165 ± 0.040 | 67.42 ± 3.58 | 86.31 ± 243.66 | 2.64 | 0.077 ± 0.025 |
| ABS Contemporary | 28 | 1.67 ± 0.11 | 1.61 | 0.846 | 0.166 ± 0.036 | 75.35 ± 2.84 | 77.63 ± 267.12 | 2.54 | 0.105 ± 0.018 |
| PLE Historic | 25 | 1.66 ± 0.06 | 1.56 | 0.939 | 0.166 ± 0.023 | 76.45 ± 5.21 | 85.67 ± 265.38 | 3.97 | 0.117 ± 0.034 |
| PLE Contemporary | 22 | 1.64 ± 0.07 | 1.49 | 1.039 | 0.164 ± 0.028 | 80.23 ± 6.64 | 91.68 ± 267.50 | 3.86 | 0.169 ± 0.031 |
| JDSP Historic | 14 | 1.69 ± 0.08 | 1.57 | 0.755 | 0.166 ± 0.032 | 124.61 ± 14.11 | 93.82 ± 287.63 | 4.50 | 0.079 ± 0.041 |
| JDSP Contemporary | 48 | 1.68 ± 0.11 | 1.54 | 1.587 | 0.183 ± 0.048 | 36.75 ± 3.73 | 102.43 ± 398.12 | 9.80 | 0.156 ± 0.046 |
| SPSP Historic | 18 | 1.75 ± 0.13 | 1.7 | 0.728 | 0.160 ± 0.055 | 111.09 ± 11.99 | 78.69 ± 336.15 | 9.44 | 0.070 ± 0.034 |
| SPSP Contemporary | 10 | 1.74 ± 0.13 | 1.55 | 0.741 | 0.149 ± 0.062 | 143.08 ± 43.76 | 102.77 ± 437.82 | 12.58 | 0.226 ± 0.077 |

**Figure 3.**
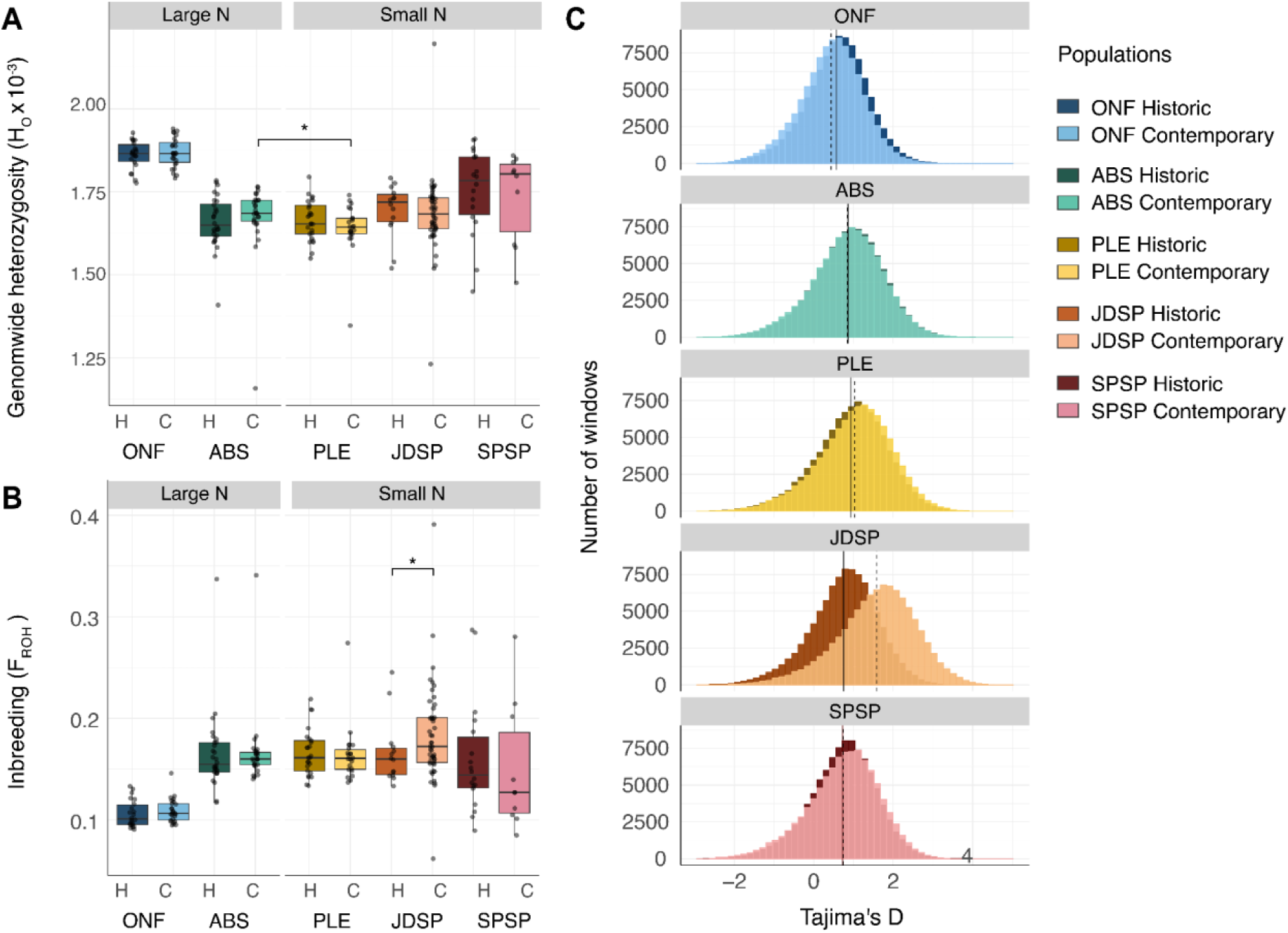
**A)** Individual genome-wide heterozygosity (*H_0_*) for each population sample. Each dot represents the total count of heterozygous nucleotide sites divided by the total nucleotide sites scored in each individual. Plotted figures appear in Table 1, column 3. **B)** Degree of inbreeding (*F_ROH_*) within the five populations. For both sampling periods, ONF had significantly higher *H_0_* and lower levels of *F_ROH_* than all other populations, even ABS. Notably, of our small populations, only the isolated population at JDSP showed a significant increase in *F_ROH_* over time. The most inbred individual belongs to the JDSP contemporary group with nearly 40% of its autosomal genome in a ROH (*F_ROH_* value of 0.39). **D)** Tajima’s *D* values in 10-kb windows. The solid, vertical black lines correspond to the mean value of the historic groups, while the dashed lines correspond to the contemporary groups. The *x-* and *y*-axes are fixed for ease of visual comparison. Positive Tajima’s *D* values indicate a deficiency of rare alleles, often associated with recent population decline, and is especially apparent in the JDSP population. For all plots, the “H” and “C” on the *x*-axis represent the historic and contemporary sampling for each population, respectively. The asterisk corresponds to *P*-values < 0.05.

Next, we compared Tajima’s *D* estimates, a sensitive indicator of recent changes in population size. We expect a recent reduction in population size results in rapid loss of rare alleles with lesser reduction in heterozygosity, driving Tajima’s *D* positive. If the decline is arrested, Tajima’s *D* will recover and even go negative as new variants persist as rare alleles (Tajima 1989). The mean Tajima’s *D* was positive among all groups, indicating a persistent population decline statewide (Figure 3C; Table 1). Across the sampling interval, Tajima’s *D* values decreased in ONF and ABS, but increased significantly in PLE, JDSP, and SPSP Contemporary, suggesting an immediate loss of rare alleles in declining populations (Table 1; Table S4). The most positive values of Tajima’s *D* were in JDSP Contemporary, followed by PLE Contemporary (1.587 and 1.039, respectively; Figure 3C; Table 1).

### Runs of homozygosity and inbreeding

In total, we detected 469,723 ROH segments in our dataset. First, we calculated the proportion of the genome in these ROH (*F_ROH_*) to investigate changes in inbreeding across sampling periods. Only the isolated population at JDSP showed a significant spike in *F_ROH_* (Wilcoxon rank-sum: *W=*436, *p=*0.047; Figure 3B; Table S5). The single most inbred individual occurred in the JDSP Contemporary group (*F_ROH_* = 0.39; Figure 3B). Interestingly, the individual with the second highest *F_ROH_* level was an outlier in our ABS Contemporary group (Figure 3B). Our detailed pedigree for this population revealed this outlier to be the progeny of a rare half-sibling mating, consistent with the unusually high *F_ROH_* for an individual in this large population.

Among historic samples, ONF individuals had significantly lower *F_ROH_* consistently across all population comparisons (Table S6). In our contemporary samples, *F_ROH_* in JDSP Contemporary was significantly higher than either ONF or ABS, and PLE harbored significantly higher *F_ROH_* than ONF (Figure 3B; Table S7). Finally, despite being our smallest population, SPSP did not have significantly higher *F_ROH_* than ONF or ABS (Table S7), consistent with known recent immigration from populations located well to the north.

Next, we investigated ROH length distributions. ONF had the lowest mean ROH length for both sampling periods, followed by ABS Contemporary and SPSP Historic (Figure 4A; Table 1). In contemporary time, SPSP had the highest number of per-individual ROH segments (143.08 ± 43.76) and longest mean ROH length across populations (102.77 ± 437.81; Figure A4), despite relatively high levels of *H_0_* compared to the other populations (Figure 3A). As expected, JDSP saw an increase in average ROH length over time (Historic: 93.82 ± 287.62 and Contemporary: 102.43 ± 398.11). ROH abundances and mean lengths were similar across the neighboring populations of ABS and PLE (Table 1).

**Figure 4.**
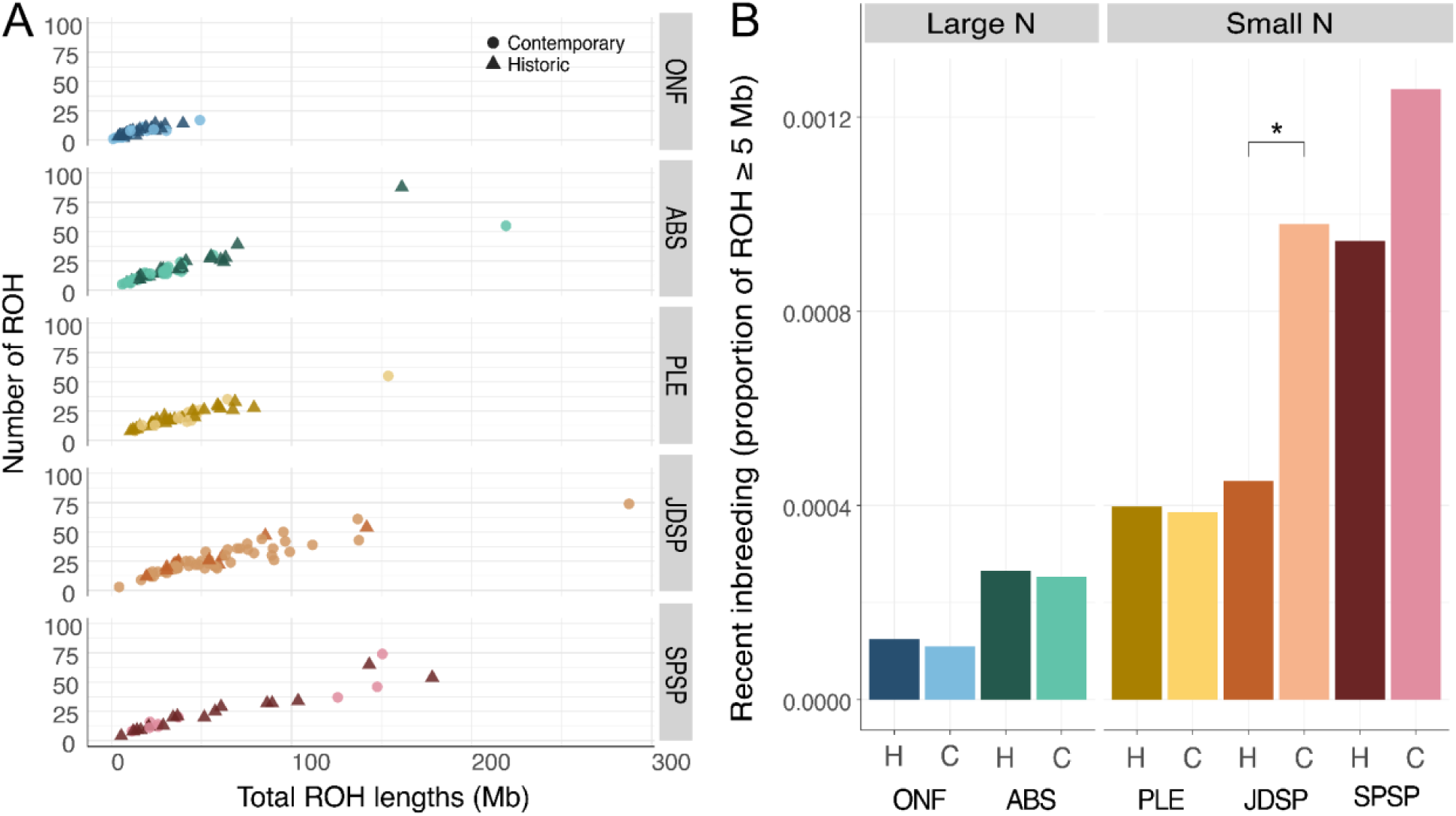
**A)** ROH abundance and lengths across sampling populations and time using SNPs without LD filtering. Each point corresponds to an individual, and the symbol (circle or triangle represent contemporary and historic timepoints, respectively). **B)** Recent inbreeding as inferred by the mean proportion of ROH ≥ 5 Mb. Only JDSP showed a significant increase in proportions across the sampling interval.

To assess recent inbreeding, we focused on the proportion of long ROH segments (≥ 5 Mb in length). Across time, we did not detect significant differences in the proportion of long ROH segments in the small, yet non-isolated PLE and SPSP populations (Figure 4B; Table S8). However, consistent with the decrease in nucleotide diversity and increase in *F_ROH_*, our isolated JDSP population showed a significant increase in the proportion of long ROH segments (Pearson’s Chi-square test: *X^2^* = 6.1806, *df* = 1, *P* = 0.0129). Finally, the large, stable ONF and ABS populations showed no significant differences in the proportion of long ROH across time (Figure 4B; Table S8).

Among populations, ONF consistently harbored a significantly lower proportion of long ROH compared to all populations except for ABS (Figure 4B; Table S8). In both sampling periods, PLE tended to have higher proportions of long ROH than ABS, although the relationship was not significant in either year (Table 1; Table S8). Interestingly, SPSP Contemporary showed the highest proportion of long ROH segments among all populations (Table S8), revealing ongoing recent inbreeding despite its relatively high genetic diversity. Finally, JDSP Contemporary had the second highest proportion of long ROH (behind SPSP Contemporary), although this relationship was not statistically significant across populations.

ROH hotspots and cold spots are of interest as they indicate regions that are unusually tolerant or intolerant of homozygosity, respectively. Applying 99th and 1st percentile cutoffs to ROH density (Figure 5, top panel), we identified 27 ROH hotspots and 31 ROH coldspots (Table S12) and identified genes that overlap these regions. The genome-wide average gene density for the FSJ genome is 14.8 genes/Mb. We found that ROH hotspots had a gene density of 13.6 genes/Mb, while coldspots had a gene density of 36.6 genes/Mb.

**Figure 5.**
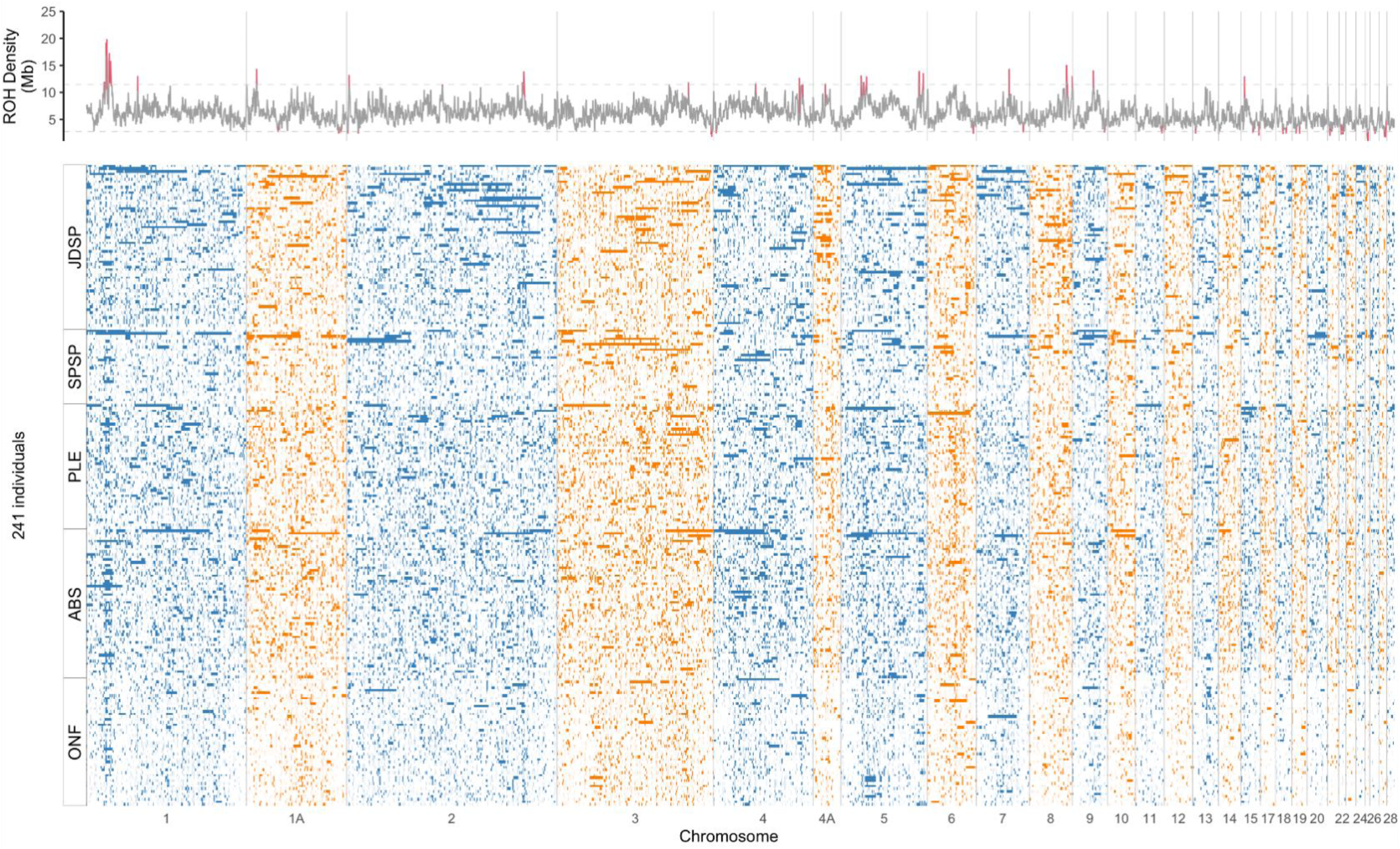
The physical mapping of ROH across all 241 individuals, grouped vertically by population. The *x*-axis represents the full span of the genome. Chromosomes are arranged left to right (alternating in blue and gold colors). Each row represents an individual. Within each population grouping, the most inbred individual, as measured by *F_ROH_*, is displayed at the top and the least inbred (lowest *F_ROH_*) is at the bottom. The density of ROH segments across all individuals is represented in the top panel as the sum of ROH segments within nonoverlapping 200-kb windows. Windows with exceptional density of ROH segments (hotspots and cold spots, defined as the top and bottom 1% respectively) are marked in red on the density plot at the top of the figure.

We performed a Gene Ontology analysis of the genes located in the ROH hotspots to identify enrichment of GO terms. Using TopGO (Alexa et al. 2023), we focused on GO terms with a TopGO Fisher p-value < 0.05. In the ROH hotspots, the Biological Process GO terms that showed an enrichment included “negative regulation of endopeptidase activity”, “negative regulation of neuroinflammatory response”, and “heterotypic cell-cell adhesion”. One possible interpretation of these regions is that they have excess ROH because of reduced genetic diversity, caused possibly by a past selective sweep, but subsequent testing of diversity in these regions found this not to be the case. With respect to Molecular Function GO terms, we found a significant enrichment of “serine-type endopeptidase inhibitor activity” and “acetylcholine receptor inhibitor activity”. We found only one GO term classified as Cellular Component that was significantly enriched: “cell surface”, a term often associated with pathogen resistance.

ROH coldspots are also of interest because they could be regions that are particularly intolerant of homozygosity, which might indicate regions that actively maintain high diversity. In ROH coldspots, the Biological Process GO terms that showed an enrichment included “anterior/posterior pattern specification”, “respiratory gaseous exchange by respiratory system”, and “tRNA pseudouridine synthesis”. These might be regions where there are recessive deleterious variants, so ROH in these regions is selectively disfavored, but we have no further evidence for this. There were no significant Molecular Function or Cellular Component GO classifications that were enriched in ROH coldspots. Table S14 presents a list of the most significant GO terms and the genes (Table S13) within the respective regions.

### Degree of relatedness and gene flow

We compared mean relatedness within populations across sampling timepoints to test our hypothesis that shrinking, isolated populations should increase in overall relatedness as they decline in size. Relatedness was quantified as the proportion of the genome of pairs of individuals that is identical-by-descent, as scored by PI_HAT (Purcell et al., 2007). Interestingly, we found a significant decrease in average pairwise relatedness in the ONF population across the sampling period, despite its relatively stable population size (Wilcoxon rank-sum test: *W* = 440, *P* = 1.47 ×10^−3^), possibly reflecting a broader sampling effort in the contemporary period or increased immigration into this large population over time. The other populations notably increased in average relatedness across time, with the largest jump within SPSP (Figure 6A; Table S9).

**Figure 6.**
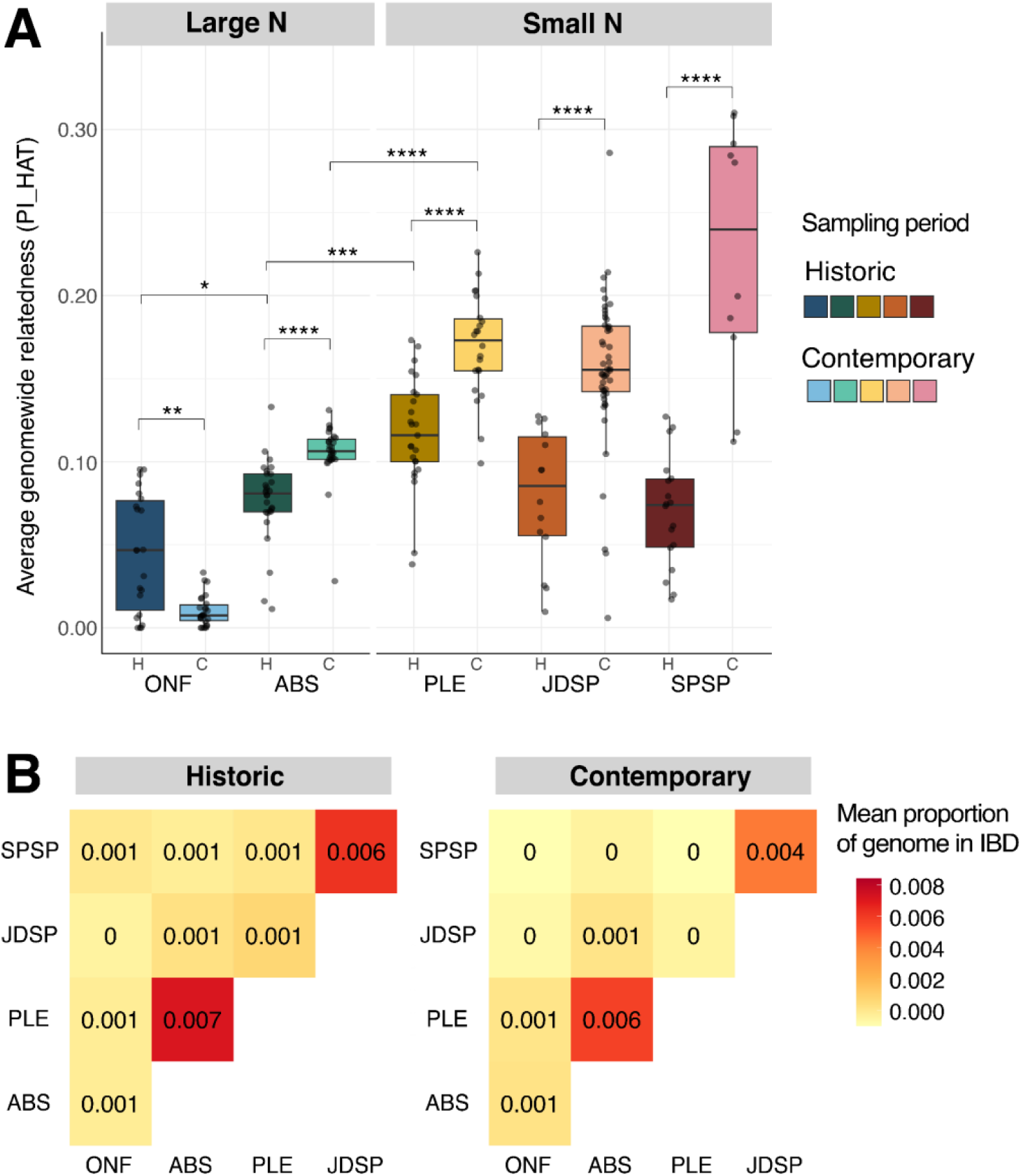
**A)** Boxplots of average within-population genome-wide relatedness (*PI_HAT*). All populations showed a significant increase in *PI_HAT* over time, except for ONF. Relatedness was higher in all declining populations compared to our large, stable populations at ONF and ABS for both sampling periods. SPSP contemporary had the widest variance, suggesting a mix of both closely and distantly-related individuals. Note that not all significant relationships are displayed in this plot. Significance is depicted as P < 0.05 (\**), P < 0.01 (**)*, P < 0.001 (***), and P < 10^−4^ (****). The “H” and “C” on the *x*-axis represent historic and contemporary sampling groups for each population. **B)** Gene flow among populations as inferred by the proportion of long IBD segments ≥ 5 Mb (within the last few generations) covering the genome. Historic (left) and contemporary (right) samples are separated. Values are rounded to the third decimal place for visualization purposes, while tiles are colored based on the continuous scale. As expected, gene flow among populations in close proximity and within the same metapopulation is elevated, with the highest gene flow found between historic JDSP and SPSP. However, IBD proportions decreased across nearly all populations in contemporary times, suggesting that statewide connectivity has been reduced (Wilcoxon rank-sum: *W*=33260346*, p*<0.001).

Next, we compared across populations within their respective sampling times to assess whether larger populations were less related, on average, than smaller populations. Interestingly, in historic samples, average relatedness in ONF was only significantly lower than ABS and PLE, but not JDSP or SPSP (Wilcoxon rank-sum test: *W =* 459, *P =* 0.033 and *W =* 34, *P =* 2.93 × 10^−6^, for ABS and PLE; Table S10). Average relatedness in historic PLE was already significantly higher than ABS and SPSP (Wilcoxon rank-sum test: *W =* 93, *P =* 1.05 × 10^−5^ and *W =* 381, *P =* 5.43 × 10^−4^, respectively; Table S10). For the contemporary samples, the large populations at ONF and ABS were both significantly less related than all three small populations (Figure 6A; Table S11). Notably, despite relatively high genetic diversity and low *F_ROH_*, relatedness values in contemporary SPSP were the highest and most varied, reflecting a mix of both highly and distantly-related individuals consistent with known occasional immigration from distant populations to the north (Table 1; Figure 6A).

Finally, we focus on long IBD segments ≥ 5 Mb to assess recent changes in gene flow among populations (within the last few generations). As expected, gene flow was the highest between ABS and PLE (within the same metapopulation) in historic times, but decreased over the sampling period (Figure 6B; proportion of long IBD = 0.007 and 0.006, respectively). Similarly, the populations of JDSP and SPSP that are in close geographic proximity but previously inferred as different metapopulations (Coulon et al. 2008), had the second highest degree of gene flow, but this proportion also decreased in contemporary times (Figure 6B; proportion of long IBD = 0.006 in historic, 0.004 in contemporary). JDSP and SPSP appeared the most isolated from the other focal populations with near zero long-IBD segments shared with ONF, ABS and PLE. Notably, overall proportions significantly decreased across nearly all populations across the sampling interval (Wilcoxon rank-sum: *W*=33260346*, p*<0.001), suggesting that connectivity is reduced statewide.

### Demographic inferences

Ancestral effective population sizes from SMC++ revealed large historic populations that all underwent steady declines over the last 100,000 years (Figure 7B). Prior to the Last Interglacial period (LIG; ∼120 kya), all FSJ populations shared similar population sizes of at least 40,000 individuals each, suggesting they may have existed as a single panmictic population. After the LIG, all populations plummeted in size up until the Last Glacial Maximum (LGM; ∼20 kya), after which there was a period of stability. By this point in FSJ evolutionary history, different demographic trajectories reflecting the current metapopulations can be observed (Figure 7B). During the period known as deglaciation (between the LGM and until the Holocene), most populations remained stable except for ONF, which saw a surprising bottleneck down to ∼5,000 individuals. At the beginning of the Holocene, ONF increased significantly to ∼10,500 individuals, recovering from the bottleneck. However, all other populations crashed during the Holocene, with JDSP suffering the greatest decline, from 10,000 to about 5,000 individuals. Finally, around 1,000 years before present population sizes once again stabilized with ONF as the largest, then SPSP, ABS, JDSP, and finally PLE at just over 4,000 individuals.

**Figure 7.**
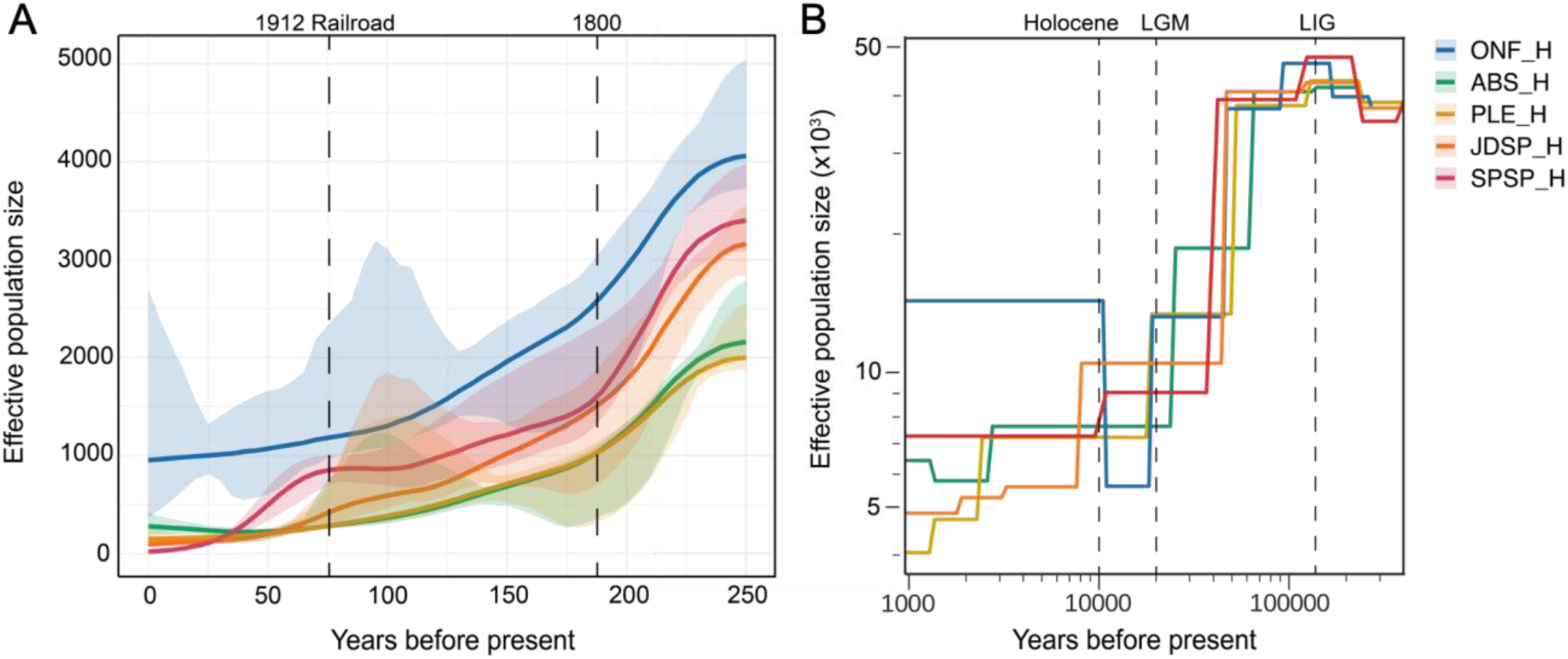
**A)** Recent and **B)** deep demographic history estimated using IBD segments or SFS, respectively. Please note the difference in scale for the effective population sizes. “H” in the legend corresponds to the 72 “historic” samples. SMC++ results in B) show a shared evolutionary history in FSJ during the LIG (Last Interglacial Period) and subsequent dramatic declines across populations leading up to the LGM (Last Glacial Maximum). Ancestral *N_e_* was high in the LIG (∼40k individuals), suggesting high connectivity and nearly panmictic gene flow during that time. Trajectories diverged as LGM approached, presumably reflecting the emergence of today’s known genetic units.

We employed a haplotype-based inference method based on IBD sharing to infer the *N_e_* across our populations within the last 50 generations and to assess the influence of anthropogenic change on FSJ demography. Starting in the late 1700s (∼45 generations ago), we saw rapid declines across all populations, with ONF remaining the largest throughout the past two centuries (Figure 7A). Interestingly, our results suggested that JDSP and SPSP were larger than either ABS or PLE until the 1900s, when they both experienced yet another catastrophic decline, coinciding with intensive human colonization following completion of the Florida East Coast Railroad (Florida Keys News, 2012). Overall, current *N_e_* estimates for each population (ONF = 952, ABS = 277, PLE = 145, JDSP = 93.9, and SPSP = 18.4) were strikingly similar to our field observations (Figure 1B) of population sizes, demonstrating the power and utility of IBD segments for population genetic analyses.

## Discussion

Understanding the genomic consequences of population decline across spatial and temporal scales is crucial for assessing extinction risks in vulnerable species. We investigated these questions in the Florida Scrub-Jay (FSJ), a well-studied system with extensive ecological and genomic resources. We sequenced whole genomes of 241 FSJs from five populations with contrasting demographic histories. We assessed genomic attributes associated with population declines and gene flow among these populations. Using genome-wide and fine-scale approaches, we reconstructed demographic histories to understand processes shaping genetic variability and lay groundwork for future evolutionary studies. Temporal sampling revealed the interplay between population size and gene flow in mediating genetic diversity and inbreeding. Sampling remnant populations allowed us to characterize genetic attributes of population loss and glimpse the potential fate of stable populations as habitat fragmentation intensifies.

Our SFS-based reconstruction showed a deep, shared demographic history across all populations that undoubtedly was shaped by the dynamic paleoclimatic history of Florida. In contrast, our haplotype-based models revealed diverging recent demographic trajectories across populations, likely brought about by accelerating anthropogenic land conversion over the past 150-200 years. Our SMC++ results suggest that during the Last Interglacial Period (∼120,000 years ago), characterized by warm and wet climate, the FSJ populations had similar and large *N_e_* (∼50 k individuals). Because this analysis assumes a closed population model, this large *N_e_* may not solely reflect population sizes but also suggest panmixia (or near-panmixia) across the five focal populations during that time (Terhorst et al., 2017). In the period leading up to the Last Glacial Maximum (∼20 kya), precipitous declines occurred across all FSJ populations as global temperatures and humidity plummeted. Although some terrestrial species likely experienced range expansions due to increasing land availability as sea levels dropped during this period (Call et al., 2016; Fehrmann et al., 2012; Guiher & Burbrink, 2008), the sharp decline in FSJ suggests that climate and its effects on local habitat availability were more important for persistence than land availability for this highly sedentary habitat specialist. Finally, as temperatures warmed again and sea levels rose leading up to the Holocene (∼10 kya), FSJ populations began to diverge in their demographic trajectories, likely reflecting the initial formation and isolation of the different genetic units and metapopulations that exist today (Stith et al. 1996, Coulon et al. 2008).

Leveraging the shared IBD segments in our sample to reconstruct the recent demographic history, our results reveal a striking portrait of the effects of anthropogenic habitat loss and urbanization on FSJ population sizes. The dramatic declines across all FSJ populations beginning in the late 18^th^ century coincide exactly with European colonization of Florida, followed by mass migration of American settlers into Florida after it became a U.S. territory in 1821 (Gregory, 2015). The citrus industry – heavily dependent on the xeric soils that harbor Florida’s oak scrub – experienced two large booms, in the late 1800s and again between the 1920s and 1980s (Florida Department of State, 2023). Our SMC++ and IBDNe results are broadly consistent in showing overall trends of declining population sizes. The SMC++ results are consistent with a fused, large panmictic population that subsequently underwent decline and fragmentation, with particularly sharp declines in JDSP and SPSP. The IBDNe analysis shows that in recent times, populations continued to decline and diverge, with ONF and ABS beginning to stabilize in size within the last 100 years. Meanwhile, the east coast populations at JDSP and SPSP experienced another notable crash in the early 1900s, ultimately dropping their *N_e_* below ABS. This decline in JDSP and SPSP coincides precisely with the completion of the southernmost stretches of the Florida East Coast Railway in 1912 (Florida Keys News, 2012). The subsequent, rapid colonization of cities and towns along the coast during the early 20th century resulted in fragmentation of the once-continuous coastal scrub. Today, JDSP represents the southernmost outpost, lacking any gene flow from once-large but now-extirpated populations that existed as far south as Miami, and cut off from northerly populations by SPSP intercepting the immigrants. Together, our demographic analyses provide the first comprehensive picture of the demographic past of the FSJ, revealing the processes that gave rise to current patterns of genomic variability.

Our genomic data confirmed previously inferred, remarkably high population structure across the relatively limited geographic distribution of the FSJ. ADMIXTURE and IBD segment analyses revealed fine-scale gene flow between certain populations consistent with their shared genetic units, as previously defined using microsatellite data (Coulon et al. 2008). Our results add to the mounting evidence that the FSJ is remarkably reluctant to disperse long distances, especially across landscapes fragmented by unsuitable habitat (Aguillon et al., 2017; Breininger, 1999; Coulon et al., 2012). Signals of ancient gene flow between central Florida (ONF and ABS) and the east coast (particularly SPSP) point to migration corridors between these historically much larger populations.Contemporary inter-population gene flow is highly reduced statewide as a result of steady increases in human land use, habitat conversion, and extensive residential development across Florida in recent decades (Boughton & Bowman, 2011). Our deep-coverage genomic dataset allowed us to document shifting gene flow among FSJ populations across space and time and revealed fine-scale admixture events that may have gone undetected without using IBD segment analysis.

Migration between PLE and the nearby population of ABS is well-documented (Nguyen et al., 2022), and both our ADMIXTURE and IBD segment analyses suggest substantial introgression of ABS ancestry into the PLE population. In SPSP where the population ranges between 15 and 30 individuals,recent field monitoring has recorded three immigration events from the north. Indeed, our SMC++ analysis shows that SPSP had the second highest *N_e_*, suggesting that this population has been well-connected throughout history and continues to receive immigrants in contemporary times despite its small population size. The proportion of shared IBD segments between SPSP and JDSP attributable to recent common descendants decreased across our sampling interval, and our ADMIXTURE results also revealed evidence of gene flow between these two populations that has essentially ceased in contemporary times. The once-substantial FSJ population south of JDSP is now extirpated, and SPSP appears as the terminus for the occasional immigrants from the north. Therefore, JDSP now appears to be fully isolated, potentially allowing for accelerated accumulation of deleterious variants.

The genomic attributes of our five focal populations were largely consistent with expectations from conservation genetics theory. As expected, the largest population (ONF) harbored higher genetic diversity (π) and lower levels of inbreeding and relatedness than any other population across both sampling timepoints. Although our intermediate-sized, stable population (ABS) saw an increase in overall relatedness between 1990 and 2017 (Fig. 3), Tajima’s *D*, which is more sensitive to allelic richness, recorded an increase in rare alleles within ABS over time. This could have arisen with the influx of PLE individuals and their unique alleles into the ABS population in contemporary times as suitable habitat disappeared in PLE.

In our small populations, we demonstrate the vital role of gene flow in mitigating the accumulation of inbreeding in small and declining populations. Despite alarmingly small population sizes (∼10 families) in the non-isolated PLE and SPSP, we found relatively low levels of inbreeding that remained unchanged across our sampling period (Figures 3 and 4). However, PLE did show significantly lower π than ABS in contemporary time, consistent with our previous studies (Nguyen et al. 2022), and Tajima’s *D* became more positive over time, indicative of the early stages of population decline. Although accumulation of ROH and the risk of recessive, deleterious variants may be staved off in these small, non-isolated populations, disappearance of unique alleles in the populations over time suggests that immigration may be waning, posing future risks to persistence of these populations.

The isolated population at JDSP displays devastating consequences of population decline. Across our 27-year sampling period, JDSP significantly increased in inbreeding and relatedness, with a higher proportion of long ROH segments derived from recent consanguineous mating. JDSP Contemporary had the highest Tajima’s *D* value out of all populations, and π also decreased across the sampling interval. JDSP further exemplifies the importance of gene flow for maintenance of genetic diversity, especially rare sequence diversity, and low levels of inbreeding. Translocation from healthy, genetically more diverse populations elsewhere across the range of the species is likely essential for small populations as isolated as JDSP is today.

ROH segments are genomic attributes that not only provide insights into the demographic histories of populations but also reveal putative regions of interest related to inbreeding depression and selection. For example, the proportion of the genome in ROH (*F_ROH_)* is a useful estimator of inbreeding in natural systems because it is correlated with homozygous mutation load (Kardos, Nietlisbach, et al., 2018; Stoffel et al., 2021b). Longer ROH are expected to harbor more deleterious alleles with larger average effects on fitness than shorter ROH where purifying selection has had more time to purge maladaptive mutations (Stoffel et al., 2021b; Szpiech et al., 2013). Furthermore, the accumulation of ROH at a specific region of the genome (known as “hotspots”) could reflect signatures of positive selection and uncover functional variants associated with inbreeding depression (Goszczynski et al., 2018; Kardos et al., 2017; Metzger et al., 2015; Zhang et al., 2015). On the other hand, regions along the genome with lower than expected abundance of ROH (“cold spots”), could indicate loci at which heterozygosity is critical for survival and thus, would be lethal or highly damaging to harbor recessive variants (Pemberton et al., 2012). Such regions are expected to undergo purging, potentially allowing for the persistence of genetically-depauperate populations with small *N_e_* (Kyriazis et al., 2023; Robinson et al., 2016, 2018). Indeed, our ROH analysis confirmed that JDSP is acquiring higher proportions of long ROH across time, indicative of recent ongoing inbreeding. SPSP also saw an increase in long ROH over time, potentially foreshadowing concerns in the future for this small population that may not be able to combat the deleterious consequences of inbreeding, even with gene flow.

Clearly, beyond effective population size, gene flow is critical in mediating levels of genetic diversity and inbreeding in fragmented species such as the FSJ. This finding is consistent with other long-term studies of wild populations where even large populations are subject to inbreeding in the absence of migration (e.g., Sattler et al., 2017). Indeed, several lines of evidence in our study and previous FSJ research (Chen et al., 2016; Summers et al., 2024) strongly suggest that gene flow even from small, genetically depauperate and inbred populations (like PLE) is critical for the persistence of seemingly stable, large subpopulations (like ABS). Even small, remnant populations should be prioritized for conservation efforts in this species.

Our population genetic estimates and inferences about the role of gene flow provide critical new data for management decisions, including identification of a candidate population (JDSP) calling for translocation and its hoped-for genetic rescue. Factors influencing population persistence, especially as it relates to genetic rescue efforts, are currently debated in the field of conservation genetics (Kardos et al., 2021; Kyriazis et al., 2021; Ralls et al., 2020; Teixeira & Huber, 2021). Namely, deleterious mutational load and the ability of small populations to purge these maladaptive variants is a complex process that is dictated by long-term demographic histories and timing, including magnitudes and duration of bottlenecks (Grossen et al., 2020; Kyriazis et al., 2021; Robinson et al., 2018). Recent studies focused on population decline in other populations, such as in the Channel Island fox (Robinson et al., 2016, 2018), Alpine ibex (Grossen et al., 2020), and Montezuma Quail (Mathur et al., 2023), suggest that populations that have experienced repeated bottlenecks and persisted at small sizes for many generations may have successfully purged deleterious recessive variation to the point that they are less susceptible to inbreeding depression (Teixeira & Huber, 2020). However, populations that have not faced recurrent bottlenecks are expected to show a strong correlation between reduced genome-wide diversity and increased inbreeding depression. For such species, maintenance of genetic diversity – including via translocation – remains a useful conservation strategy (Kardos et al., 2021; Ralls et al., 2020). Our detailed demographic inferences and genomic characterization of focal FSJ populations presents a foundation for forthcoming investigations into the deleterious mutational load and precise population genetic simulations within this species. Our study exemplifies how consideration of all aspects of a population’s genetic attributes, from the degree of isolation to detailed demographic history to fine-scale genomic attributes, should be evaluated for a comprehensive understanding of how genomic dynamics can contribute to conservation decisions.

## Conclusions

Our comprehensive genomic analysis of five FSJ populations provides important details about demographic histories, genetic diversity, and the impact of anthropogenic activities on this vulnerable species. Our data underscore the critical role of gene flow in maintaining genetic diversity and mitigating inbreeding depression, even in small and fragmented populations. The populations at ONF and ABS have stabilized in recent times, while only the isolated population JDSP (as opposed to non-isolated PLE and SPSP) decreased in nucleotide diversity and accumulated ROH segments possibly harboring deleterious variants. These results point to the importance of both conserving small, remnant, “stepping stone” populations and, in the extreme cases, facilitating gene flow through translocation programs, especially given the evidence of ongoing, statewide reduction in connectivity. Our research provides a critical evaluation of a rapidly declining species, offering immediate insights for conservation management, and establishes essential genomic resources and a foundation for future studies on fundamental evolutionary and population genetic processes in contemporary times.

## Lead contact

Further information and queries should be directed to and will be fulfilled by the lead contact, Tram Nguyen

## Materials availability

There are no newly generated materials associated with this study.

## Data and code availability

- Raw whole-genome sequences for this study will be uploaded to NCBI upon submission of this paper.
- The variant calling pipeline and code for performing the analyses in this manuscript are available at https://github.com/tram-nguyen-n/FSJ_WGS20x.
- Any additional information required to reanalyse the data reported in this paper is available from the lead contact upon request.

## Acknowledgments

Co-author Reed Bowman (1958–2023) was a fun, inspirational colleague and an invaluable source of FSJ knowledge. We thank Rob Rossmanith and Angela Tringali for providing field supplies and knowledge, as well as the JayWatch volunteers who assisted with sample collection and demographic monitoring, with special thanks to Liz Hailman. We’d like to thank Bronwyn Butcher and Asha Jain for their laboratory preparation advice and help. Thank you to the Cornell BioHPC and Statistical Consulting Unit (CSCU) for their computational and statistical guidance, and Tyler Litheroth for his guidance on the demographic modeling.

Finally, thank you to Ariel Fogel for providing feedback on the manuscript and Faye Romero helping with queries about the linkage map.

## Funding

T.N.N. was supported by the NSF Graduate Fellowship Program and by the Cornell Lab of Ornithology (Gomez Fund). The following grants supported this project: Cornell Lab of Ornithology Athena Fund, Florida Ornithological Society Cruickshank Award, American Ornithological Society Mewaldt-King Research Award, Cornell Sigma Xi Research Award, and Cornell University, Andrew W. Mellon Student Research Grant.

## Author contributions

Designed project: T.N.N., A.G.C., and J.W.F. Performed field-work and sample collection: T.N.N, N.L., R.B., K.M., and J.W.F. Assisted with funding for sample collection: J.W.F. and S.F. Obtained genomic data: T.N.N. Primary genomic analysis T.N.N., E.J.C., and N.C., with contribution from M.L. and F.E.B. Provided methodological or experimental guidance: A.G.C., E.J.C., N.C., S.F., M.L., F.E.B, and J.W.F. Supervised research: A.G.C. and J.W.F. Wrote manuscript: T.N.N., E.J.C., A.G.C., J.W.F., with edits from all authors.

## Declaration of interests

The authors declare no competing interests.

## Supporting Information

Supplementary files are available for download.

## STAR★Methods

### Software Resources Table

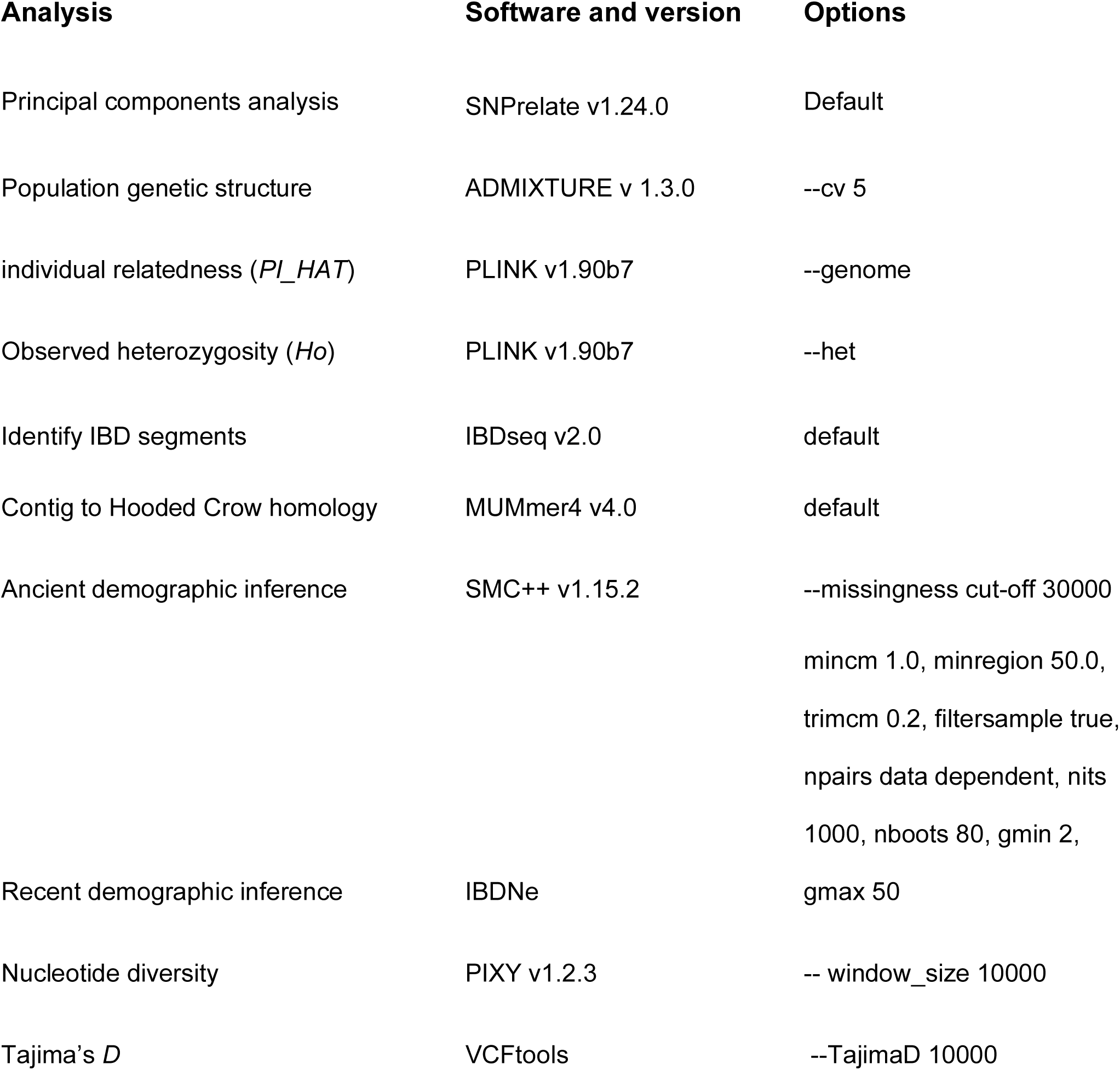
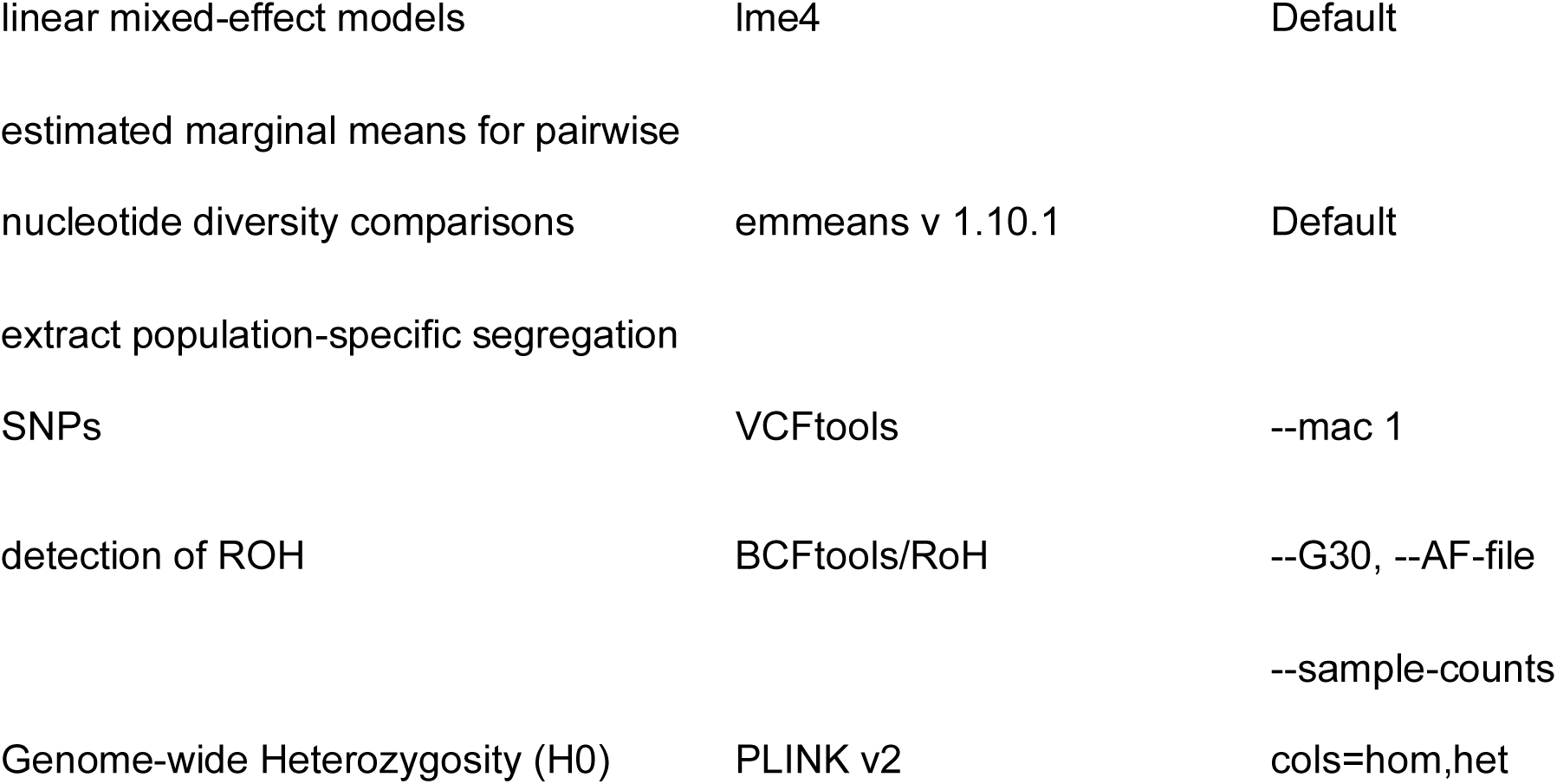

### Methods

#### Sampling and sequencing

We collected a total of 241 blood samples from breeding individuals from our five focal populations across two timepoints. We obtained 107 historic blood samples collected between the years 1990-1993 for four populations ONF (*n*=22), ABS (*n*=28), PLE (*n*=25), JDSP (*n*=14) and from the year 2004 for SPSP (*n*=18). Contemporary blood samples for 134 individuals were collected in the years 2017-2019 for each population: ONF (*n*=26), ABS (*n*=28), PLE (*n*=22), JDSP (*n*=48), and SPSP (*n*=10). Blood was sampled from the brachial vein in each bird’s wing via a needle prick and stored in lysis buffer at room temperature. We limited our genomic analyses to jays known to be breeding adults to avoid including closely related individuals (i.e., within the same nuclear family) which could bias our genetic estimates. DNA was extracted using the DNEasy blood and tissue kit (Qiagen, CA, USA) and sent to Novogene (Davis, California) for library preparation and 2×150 paired-end whole-genome resequencing on the Illumina NovaSeq and HiSeq platforms (Illumina, CA). Mean sequencing coverage was 20.7x and ranged from 17.2x to 50.1x (this latter JDSP individual was sequenced twice to achieve adequate coverage).

#### Read alignment and variant calling

We assessed sequencing read quality using FastQC v 0.11.8 (Simon, 2010), trimmed raw reads for adaptor sequences, and applied the following quality filters using Trimmomatic v 0.36 (Bolger et al., 2014). Following GATK’s best practices pipeline for germline variant detection (Van der Auwera et al., 2013), we mapped reads using the aligner BWA MEM v 0.7.13 (Li & Durbin, 2009) to a Florida Scrub-Jay reference genome (Driscoll et al. 2024) created with a combination of Illumina short-reads, Hi-C, and Chicago reads (Elbers et al., 2019). PCR duplicates were marked with Picard Tools v 2.8.2 and genotypes were called using HaplotypeCaller in GATK v3.8 (Poplin et al., 2018; Van der Auwera et al., 2013) resulting in one file containing all sites along the genome. Variants were selected with SelectVariants and filtered for quality with the VariantFiltration tool (DePristo et al., 2011; Van der Auwera et al., 2013). Variants passing GATK’s hard filtering then underwent *post hoc* filtering for genotype quality of greater than 20 and minimum site quality of greater than 30 in VCFtools (Danecek et al., 2011). At this point, samples with more than 20% missing sites after our quality filtering were excluded from the study, yielding the final 241 samples to be analyzed (out of 291 original samples). Sites that had less than half or more than double the average depth were also removed with BCFtools (Danecek et al., 2021). We retained only biallelic single-nucleotide polymorphic sites (SNPs) that were missing in fewer than 15% of our samples (F_MISSING > 0.15 in BCFtools) and excluded Z-linked SNPs. We used our two largest and most stable populations (ONF and ABS, combined across sampling years) to test all SNPs for Hardy-Weinberg equilibrium. Autosomal variants with *P-*values < 1 × 10^−6^ for Hardy-Weinberg equilibrium were removed. We produced two final datasets for downstream analyses: one with and one without linkage disequilibrium pruning (8,901,142 or 4,555,333 biallelic SNPs, respectively, across the 241 breeders).

To generate a dataset of high-confident invariant and variant sites together, we followed the recommended workflow by Korunes & Samuk (2021). After calling haplotypes in HaplotypeCaller, we generated a file containing both variant and invariant sites using GenotypeGVCFs (--all-sites). As recommended by Korunes & Samuk (2021), we applied the GATK hard filters and additional post hoc filtering in VCFtools as previously described to both our variant and invariant data separately, and omitted Hardy-Weinberg filtering on invariant sites which resulted in 912,103,392 invariant sites of high confidence. We combined these with our high-confidence 8,901,142 bi-allelic SNPs in Hardy-Weinberg equilibrium for a total of 921,004,534 non-missing sites. This dataset was used to calculate π and Tajima’s *D*.

#### Population structure and genetic ancestry

We first assessed broad-scale genetic structure across all populations and timepoints using a principal components analysis (PCA; Figure S1) with the R package SNPrelate v1.24.0 (Zheng et al., 2012). To determine fine-scale genetic structure across population groups and timepoints, we used the program ADMIXTURE v1.3.0 (Alexander et al., 2009) to estimate the ancestral composition of our samples. We ran ADMIXTURE for a range of *K* clusters from 1 to 10 and calculated the cross-validation error across 5 repetitions for each *K* value. We selected *K*=7 as the optimal number of clusters given that yielded the lowest cross-validation error, with *K*=4 as the second most optimal cluster number, consistent with our broad expectations of four metapopulations identified by Coulon et al. (2008).

#### Genetic diversity

Using our dataset of all confidently-called invariant and biallelic variant sites, we calculated genome-wide observed heterozygosity (*H_O_*) at the nucleotide level for each individual with Plink2 (Chang et al. 2015). *H_O_* was calculated as the proportion of observed heterozygosity sites out of all confidentially-called heterozygosity and invariant sites per individual genome. We estimated nucleotide diversity (π) in 10 kb windows with PIXY v1.2.3 (Korunes & Samuk, 2021). We then applied a Wilcoxon rank-sum test to further compare *H_O_* between the historic and contemporary samples within populations and across population groups, corrected for multiple-testing with the Bonferroni method.

We estimated Tajima’s *D* in 10-kb windows with VCFtools (--TajimaD 10000). To statistically compare Tajima’s *D* between populations and across sampling periods, we fitted a linear mixed-effect model using the R package lme4 (Bates et al. 2015), setting Tajima’s *D* as the response variable and population, sampling time, the interaction between the two as fixed effects. We set the chromosome as a random effect to account for any correlation at the chromosome level between our response variables. Using our linear model, we then made post hoc pairwise comparisons between groups and sampling times by calculating estimated marginal means (EMM) with the R package emmeans (Lenth et al. 2018). All *P*-values were adjusted for multiple testing using the Tukey method.

#### Runs-of-homozygosity and inbreeding

We detected ROH using BCFtools/RoH which utilizes a hidden Markov model to discriminate between autozygous and non-autozygous regions while incorporating information on allele frequency (Narasimhan et al., 2016). We used our dataset of high-confidence SNPs and omitted LD pruning for this analysis as it often reduces detection of ROH and thus biases estimates of *F_ROH_* (Meyermans et al., 2020). Because the number of polymorphic sites varies across populations, we extracted SNPs that were segregating within each population using VCFtools and ran BCFtools/RoH on each population separately. However, we used allele frequencies derived from our entire dataset of 8,901,142 SNPs across 241 breeders with the command for more accurate expected allele counts, especially given our small sample sizes for populations such as JDSP Historic (*n*=12) and SPSP Contemporary (*n*=9). We retained ROH segments that contained a minimum of 5 SNPs and had posterior quality scores ≥ 15. We report the mean number of ROH segments per individual (to account for differences in sampling sizes) and mean ROH lengths for each population.

To estimate levels of inbreeding for each individual, we calculated the proportion of the genome in runs-of-homozygosity (*F_ROH_*). To determine if inbreeding increased with population decline, we compared *F_ROH_* across time periods for each population. Because we expected our large, intensively-managed populations at ONF and ABS to remain stable, we applied a two-tailed Wilcoxon rank-sum test to compare the difference in mean inbreeding between the historic and contemporary samples for these two populations. As we expected inbreeding to increase over time in our small populations of PLE, JDSP, and SPSP, we applied a one-tailed Wilcoxon rank-sum test to assess the directionality of *F_ROH_* across our temporal samples.

We also estimated the number and length distribution of ROH segments across the genome. Focusing on the longest ROH, those ≥ 5 Mb in length (approximately 30.3 cM based on our sex-averaged linkage map; Romero et al. 2024), arising from inbreeding within the last few generations, we tested for significant differences in the proportion of long ROH segments across years and populations using a Pearson’s Chi-Square test of proportions to assess whether declining populations showed evidence of recent inbreeding.

Finally, we investigated regions of the genome with unusually high (“hotspots”) or low (“cold spots”) abundance of ROH. Establishing the statistical significance of these regions involves detailed knowledge of the linkage map as well as estimates of population parameters like the rates of mutation and migration. We instead focused on the top and bottom 1% in the distribution of ROH across the genome.

#### Gene ontology enrichment analysis

For each ROH hotspot and coldspot identified as falling in the top and bottom 1%, we extracted the genes within these regions and investigated their biological functions using a gene ontology (GO) enrichment approach. We mapped the identified ROH regions to the most recent FSJ genome where the annotations have been curated (Romero et al. 2024). We required that more than 50% of the region mapped with greater than 98% identity. Five coldspot regions did not map using these thresholds, as shown in Table S12. Using the mapped coordinates, we identified genes in each region and the corresponding GO terms from the most recent FSJ genome annotation (Romero et al. 2024). Finally, using the topGO v2.42.0 package in R (Alexa & Rahnenfuhrer, 2023), we conducted GO term enrichment analyses for genes within the ROH hotspots and coldspots and tested for GO term enrichment with a Fisher’s Exact Test using a *P*-value threshold of 0.05.

#### Degree of relatedness and gene flow

To assess the degree of relatedness across sampling periods and between populations, we calculated the proportion of the genome that is identical-by-descent (IBD) across all pairs of individuals (*PI_HAT*) using PLINK (Purcell et al., 2007). We compared mean *PI_HAT* within populations across sampling timepoints using a Wilcoxon sum rank test to determine whether shrinking, isolated populations showed increased relatedness as they declined in size. We also compared *PI_HAT* across populations to assess whether individuals from small populations were more related, on average, than individuals in larger populations. We also inferred segments that were identical due to shared common ancestry (IBD segments) between individuals across populations and assessed their length distributions as a metric of gene flow between populations. To accurately identify IBD segments, we used the program IBDseq (Browning & Browning, 2013). Unlike our previous analysis in PLINK which reports the proportion of the genome that is shared between pairs of individuals within the same population, this analysis focuses specifically on very long IBD segments (i.e., ≥ 5 Mb). We selected this threshold given that we can reliably detect longer IBD segments > 1 Mb, and while arbitrary, segments ≥ 5 Mb truly reflect long IBD occurring from recent shared parentage. To capture these long IBD segments, we omitted LD pruning and restricted our analysis to the 29 largest and most complete contigs, which had inferred homology to the Zebra Finch using the program MUMmer4 (Marçais et al., 2018; Weissensteiner, 2020). Because these long, identical segments of DNA cannot have arisen by any means other than recent shared ancestry, they harbor reliable information about connectivity between populations. To assess this metric, we compared the proportion of long IBD segments shared between populations, normalized for different sample sizes. This proportion was calculated as the count of IBD segments ≥ 5 Mb divided by the number of pairs of individuals compared.

#### Demographic inference

To investigate ancient changes in population sizes in our populations (>1k years in the past), we used the program SMC++ v1.15.2 (Terhorst et al., 2017). We used putatively neutral sites from unrelated individuals in our historic sample, collected before recent suspected bottlenecks, so as not to bias our estimates with long ROH. Specifically, we filtered our dataset of high-confidence autosomal SNPs to exclude genic regions and removed highly related individuals with a pairwise genome-wide IBD of > 0.2, as estimated from PLINK. Because SMC++ utilizes LD information in a coalescent hidden Markov model, we omitted LD filtering for this analysis. Finally, we used only variants within our 29 chromosome-level contigs that correspond to Zebra Finch chromosomes (Weissensteiner et al., 2017). Our resulting dataset includes a total of 72 individuals from the historic groups (ONF: *n*=11, ABS: *n*=24, PLE: *n*=16, JDSP: *n*=9, and SPSP: *n*=12) and 4,419,566 biallelic SNPs in non-coding regions. We then randomly selected five individuals from each population to create a composite likelihood per scaffold to vary the identity of the “distinguished” individuals to which all other samples were compared (following Walsh et al., 2021; Terhorst et al., 2017). To remove false signals produced by recent bottlenecks, we disregarded contiguous stretches of homozygosity greater than 30 kb and estimated historical population sizes across 20 independent replicate analyses for more reliable estimates of *N_e_*. For these demographic analyses, we used the collared flycatcher mutation rate of 2.3 × 10^−9^ per year (Smeds et al., 2016) and a five year generation time (Nguyen et al., 2022).

Finally, given that the effective population size (*N_e_*) affects the probability of randomly drawing two alleles that are IBD (e.g., smaller *N_e_* results in a higher chance of randomly drawing a shared allele), we leveraged the IBD segments from our 72 historic samples previously mentioned to reconstruct the demographic histories across our populations by estimating a time series of effective population sizes in the recent past (up to 50 generations). We implemented the analysis in the program IBDNe (Browning & Browning 2015), aided also by our sex-averaged FSJ recombination map estimated from nearly 4,000 pedigreed individuals (Romero et al. 2024). These haplotype-based methods contrast with conventional likelihood-based models which focus on the site frequency spectra (SFS), rather than patterns of LD, to capture coalescent events further in the past, and thus recapitulate deeper demographic history (Excoffier et al., 2013; Gutenkunst et al., 2009; Li & Durbin, 2011; Mather et al., 2020; Terhorst et al., 2017; Terhorst & Song, 2015). Given the dramatic increase in the local human population, housing development, and industrialization in Florida around the 1800s (approximately 250 years ago, equivalent to 50 FSJ generations), we used IBDNe to assess the consequences of relatively recent anthropogenic change on FSJ population sizes.

## Notes

### Competing Interest Statement

The authors have declared no competing interest.

