## Supplemental Figures S1-S2 and Tables S1-S14 for "Whole-genome sequencing across space and time reveals impact of population decline and reduced gene flow in Florida Scrub-Jays"

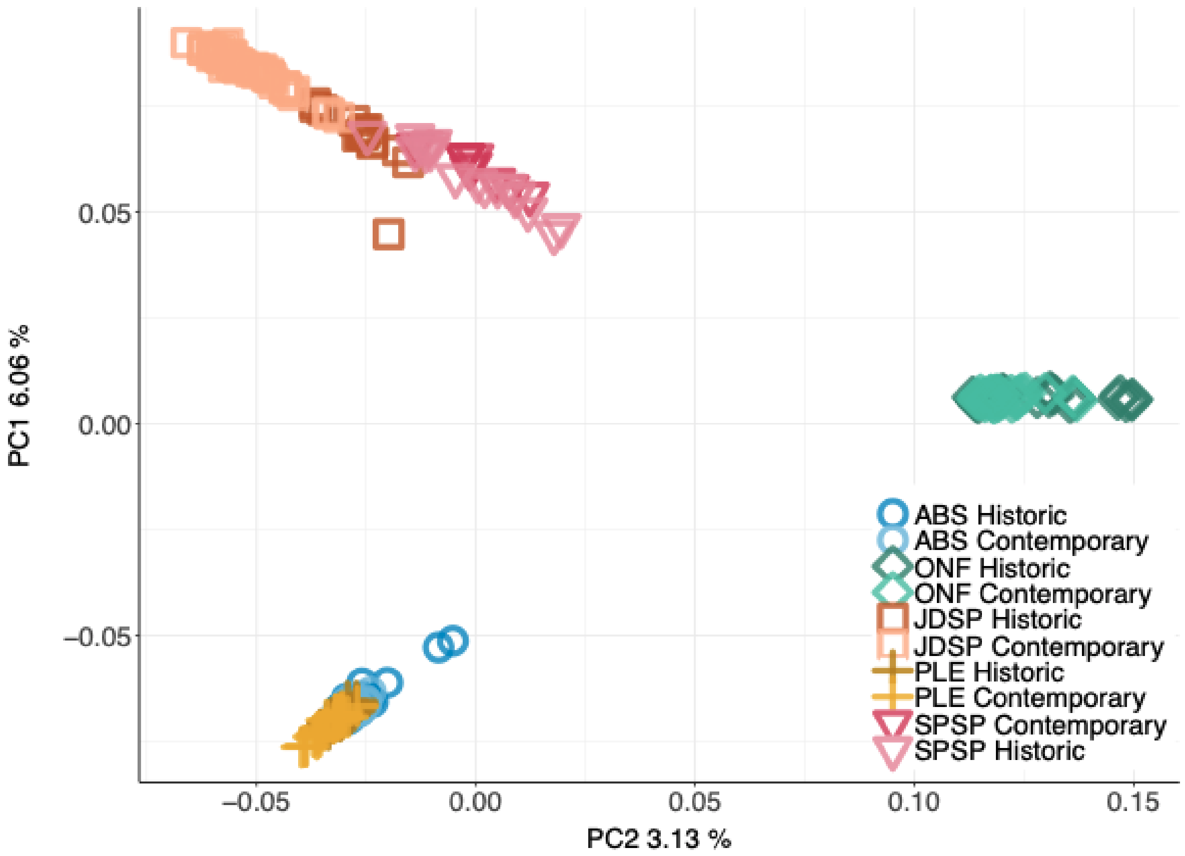


**Figure S1.** Principal components analysis (PCA) using 8,901,142 autosomal, biallelic SNPs of high-confidence and quality. Each point represents a breeding individual included in the main study (n=241). Genetic clustering determined by the PCA is consistent with the metapopulation structure detected in previous studies using microsatellite data (Coulon et al., 2008; Stith et al., 1996).

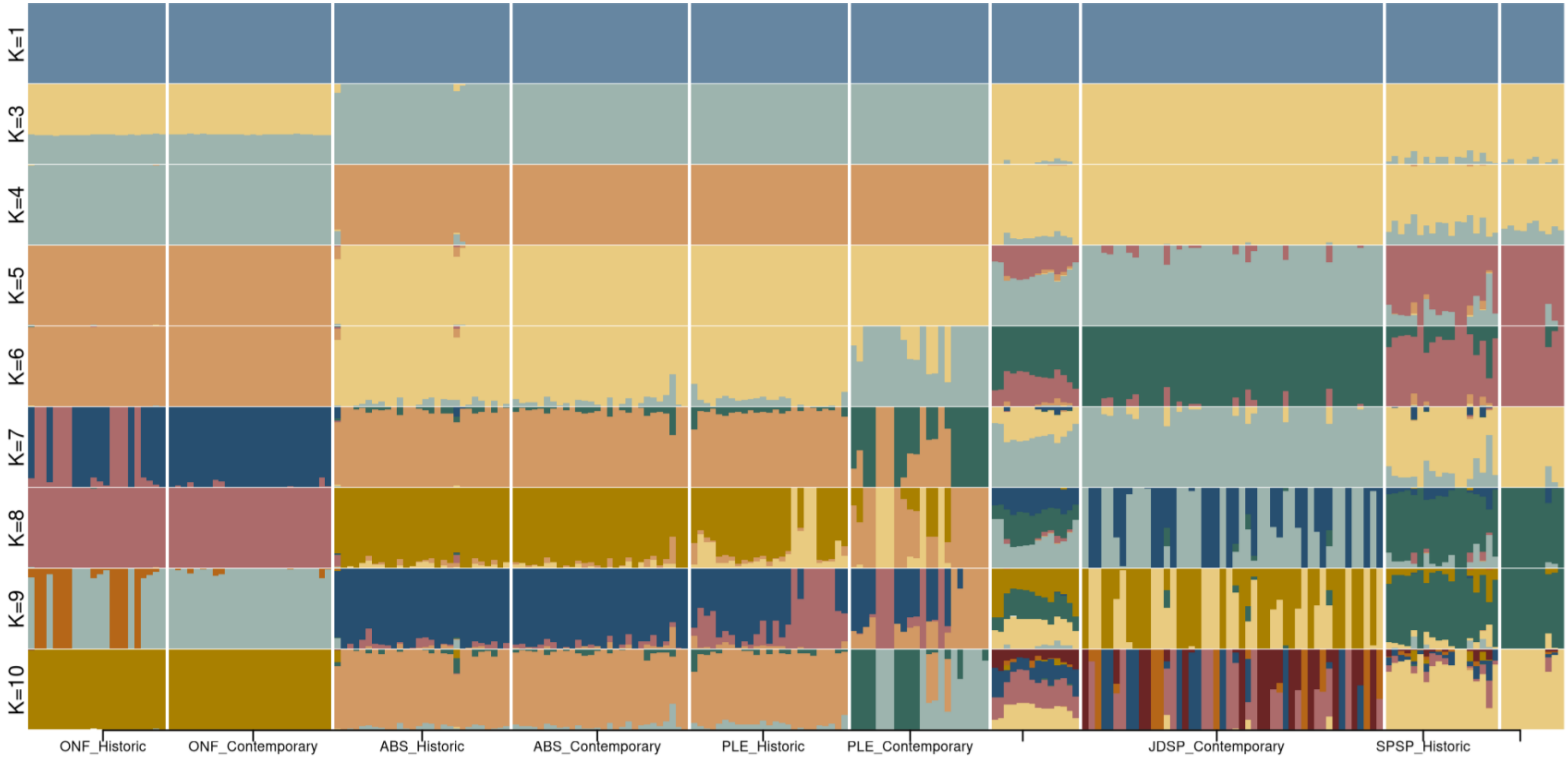


**Figure S2.** ADMIXTURE plots showing the composition of genetic ancestry for all individuals across multiple genetic clusters (*K*=1 to 10). Each column is an individual with the proportion of each color showing the proportion of ancestry assigned to a genetic group. We selected *K*=7 as the optimal number of clusters given that yielded the lowest cross-validation error, with *K*=4 as the second most optimal cluster number, consistent with our broad expectations of the 4 metapopulations.

**Table S1.** Test for significant differences in mean heterozygosity ($H_o$) within populations across the sampling interval using Wilcoxon rank sum test.

| Population Comparison | Test Statistic (W) | p-value |
|---|---|---|
| ONF Historic - ONF Contemporary | 197 | 0.0667 |
| **ABS Historic - ABS Contemporary** | **189** | **0.0007** |
| PLE Historic - PLE Contemporary | 293 | 0.6520 |
| **JDSP Historic - JDSP Contemporary** | **177** | **0.0033** |
| SPSP Historic - SPSP Contemporary | 76 | 0.2620 |

**Table S2.** Test for significant differences in mean heterozygosity ($H_o$) between historic populations groups using Wilcoxon rank sum two-tailed test, adjusted for multiple testing.

| Population Comparison | Test Statistic (W) | p-value | Bonferroni-adjusted p-value |
|---|---|---|---|
| **ONF Historic - PLE Historic** | **550** | **1.35E-13** | **1.35E-12** |
| **ONF Historic - ABS Historic** | **3** | **1.58E-13** | **1.58E-12** |
| **ONF Historic - SPSP Historic** | **396** | **1.76E-11** | **1.76E-10** |
| **ONF Historic - JDSP Historic** | **0** | **5.27E-10** | **5.27E-09** |
| **JDSP Historic - PLE Historic** | **286** | **7.71E-04** | **8.00E-03** |
| **ABS Historic - PLE Historic** | **529** | **1.00E-03** | **1.10E-02** |
| PLE Historic - SPSP Historic | 129 | 1.80E-02 | 1.76E-01 |
| JDSP Historic - SPSP Historic | 104 | 4.19E-01 | 1.00E+00 |
| ABS Historic - JDSP Historic | 161 | 3.62E-01 | 1.00E+00 |
| ABS Historic - SPSP Historic | 204 | 2.88E-01 | 1.00E+00 |

**Table S3.** One-tailed Wilcoxon rank sum test to statistically test the hypothesis that larger populations (ONF and ABS) have higher $H_o$ than small populations (PLE, JDSP, SPSP) in contemporary samples. Tests are adjusted for multiple testing.

| Population Comparison | Test Statistic (W) | p-value | Bonferroni-adjusted p-value |
|---|---|---|---|
| **ONF Contemporary > JDSP Contemporary** | **26** | **2.20E-16** | **6.60E-16** |
| **ONF Contemporary > SPSP Contemporary** | **1** | **7.87E-09** | **2.36E-08** |
| **ONF Contemporary > PLE Contemporary** | **0** | **3.65E-14** | **1.10E-13** |
| ABS Contemporary > JDSP Contemporary | 732 | 7.41E-01 | 1.00E+00 |
| ABS Contemporary > SPSP Contemporary | 200 | 9.78E-01 | 1.00E+00 |
| ABS Contemporary > PLE Contemporary | 312 | 5.35E-01 | 1.00E+00 |

**Table S4.** T-test for significance between Tajima's D across populations for historic and contemporary groups after fitting our linear model. P-values are adjusted for multiple testing.

Historic samples

| Contrast | Estimate | SE | df | t.ratio | p-value |
|---|---|---|---|---|---|
| **ABS - JDSP** | **0.09607** | **0.0166** | **261** | **5.798** | **<.0001** |
| **ABS - ONF** | **0.26058** | **0.0166** | **261** | **15.726** | **<.0001** |
| **ABS - PLE** | **-0.08307** | **0.0166** | **261** | **-5.013** | **<.0001** |
| **ABS - SPSP** | **0.10442** | **0.0166** | **261** | **6.302** | **<.0001** |
| **JDSP - ONF** | **0.16451** | **0.0166** | **261** | **9.928** | **<.0001** |
| **JDSP - PLE** | **-0.17914** | **0.0166** | **261** | **-10.811** | **<.0001** |
| JDSP - SPSP | 0.00835 | 0.0166 | 261 | 0.504 | 0.9869 |
| **ONF - PLE** | **-0.34365** | **0.0166** | **261** | **-20.739** | **<.0001** |
| **ONF - SPSP** | **-0.15616** | **0.0166** | **261** | **-9.424** | **<.0001** |
| **PLE - SPSP** | **0.18749** | **0.0166** | **261** | **11.315** | **<.0001** |

Contemporary samples

| Contrast | Estimate | SE | df | t.ratio | p-value |
|---|---|---|---|---|---|
| **ABS - JDSP** | **-0.74421** | **0.0166** | **261** | **-44.912** | **<.0001** |
| **ABS - ONF** | **0.37829** | **0.0166** | **261** | **22.83** | **<.0001** |
| **ABS - PLE** | **-0.20291** | **0.0166** | **261** | **-12.245** | **<.0001** |
| **ABS - SPSP** | **0.06715** | **0.0166** | **261** | **4.052** | **0.0006** |
| **JDSP - ONF** | **1.1225** | **0.0166** | **261** | **67.742** | **<.0001** |
| **JDSP - PLE** | **0.5413** | **0.0166** | **261** | **32.667** | **<.0001** |
| **JDSP - SPSP** | **0.81136** | **0.0166** | **261** | **48.965** | **<.0001** |
| **ONF - PLE** | **-0.5812** | **0.0166** | **261** | **-35.075** | **<.0001** |
| **ONF - SPSP** | **-0.31115** | **0.0166** | **261** | **-18.777** | **<.0001** |
| **PLE - SPSP** | **0.27005** | **0.0166** | **261** | **16.297** | **<.0001** |

**Table S5.** Test for significant differences in mean inbreeding ($F_{ROH}$) within populations across the sampling interval using Wilcoxon rank sum test.

| Population Comparison | Test Statistic (W) | P-value |
|---|---|---|
| ONF Historic - ONF Contemporary | 343 | 0.245 |
| ABS Historic - ABS Contemporary | 419 | 0.667 |
| PLE Historic - PLE Contemporary | 250 | 0.705 |
| **JDSP Historic - JDSP Contemporary** | **436** | **0.047** |
| SPSP Historic - SPSP Contemporary | 69 | 0.846 |

**Table S6.** Test for significant differences in mean inbreeding ($F_{ROH}$) between historic populations groups using Wilcoxon rank sum two-tailed test, adjusted for multiple testing.

| Population Comparison | Test Statistic (W) | p-value | Bonferroni-adjusted p-value |
|---|---|---|---|
| **ONF Historic - PLE Historic** | **0** | **1.35E-13** | **1.35E-12** |
| **ONF Historic - ABS Historic** | **606** | **3.13E-12** | **3.13E-11** |
| **ONF Historic - JDSP Historic** | **308** | **5.27E-10** | **5.27E-09** |
| **ONF Historic - SPSP Historic** | **47** | **1.15E-05** | **1.15E-04** |

**Table S7.** One-tailed Wilcoxon rank sum test to statistically test the hypothesis that larger populations (ONF and ABS) have lower inbreeding ($F_{ROH}$) than small populations (PLE, JDSP, SPSP) in contemporary samples. Tests are adjusted for multiple testing.

| Population Comparison | Test Statistic (W) | p-value | Bonferroni-adjusted p-value |
|---|---|---|---|
| **ONF Contemporary < JDSP Contemporary** | **1217** | **5.37E-16** | **1.61E-15** |
| ONF Contemporary < SPSP Contemporary | 73 | 2.24E-02 | 6.71E-02 |
| **ONF Contemporary < PLE Contemporary** | **4** | **4.38E-13** | **1.31E-12** |
| **ABS Contemporary < JDSP Contemporary** | **458** | **1.04E-02** | **3.13E-02** |
| ABS Contemporary < SPSP Contemporary | 199 | 9.76E-01 | 1.00E+00 |
| ABS Contemporary < PLE Contemporary | 316 | 5.65E-01 | 1.00E+00 |

**Table S8.** Chi-square test of proportions for long ROH segments among historic populations.

| Population Comparison | Chi-square statistics | Bonferroni-adjusted p-value |
|---|---|---|
| ONF Historic - ABS Historic | X-squared = 2.5226, df = 1, p-value = 0.1122 | 1.1220 |
| ONF Historic - PLE Historic | **X-squared = 6.898, df = 1, p-value = 0.008629** | 0.0863 |
| ONF Historic - JDSP Historic | **X-squared = 7.3783, df = 1, p-value = 0.006601** | 0.0660 |
| ONF Historic - SPSP Historic | **X-squared = 29.252, df = 1, p-value = 6.354e-08** | 0.0000 |
| PLE Historic - SPSP Historic | **X-squared = 9.7151, df = 1, p-value = 0.001828** | 0.0183 |
| ABS Historic - JDSP Historic | X-squared = 1.7778, df = 1, p-value = 0.1824 | 1.8240 |
| ABS Historic - PLE Historic | X-squared = 1.3499, df = 1, p-value = 0.2453 | 2.4530 |
| ABS Historic - SPSP Historic | **X-squared = 18.326, df = 1, p-value = 1.862e-05** | 0.0002 |
| JDSP Historic - PLE Historic | X-squared = 0.1082, df = 1, p-value = 0.7422 | 7.4220 |
| JDSP Historic - SPSP Historic | **X-squared = 4.7761, df = 1, p-value = 0.02886** | 0.2886 |

**Table S9.** Chi-square test of proportions for long ROH segments among contemporary populations

| Population Comparison | Chi-square statistics | Bonferroni-adjusted p-value |
|---|---|---|
| ONF Contemporary - JDSP Contemporary | **X-squared = 7.3783, df = 1, p-value = 0.006601** | 0.0066 |
| ONF Contemporary - ABS Contemporary | X-squared = 2.5226, df = 1, p-value = 0.1122 | 0.1122 |
| ONF Contemporary - PLE Contemporary | **X-squared = 6.898, df = 1, p-value = 0.008629** | 0.0086 |
| ONF Contemporary - SPSP Contemporary | **X-squared = 29.252, df = 1, p-value = 6.354e-08** | 0.0000 |
| JDSP Contemporary - ABS Contemporary | X-squared = 1.7778, df = 1, p-value = 0.1824 | 0.1824 |
| JDSP Contemporary - SPSP Contemporary | **X-squared = 4.7761, df = 1, p-value = 0.02886** | 0.0289 |
| JDSP Contemporary - PLE Contemporary | X-squared = 0.1082, df = 1, p-value = 0.7422 | 0.7422 |
| ABS Contemporary - PLE Contemporary | X-squared = 1.3499, df = 1, p-value = 0.2453 | 0.2453 |
| ABS Contemporary - SPSP Contemporary | **X-squared = 18.326, df = 1, p-value = 1.862e-05** | 0.0000 |
| PLE Contemporary - SPSP Contemporary | **X-squared = 9.7151, df = 1, p-value = 0.001828** | 0.0018 |

**Table S10.** Test for significant differences in mean relatedness (*PI_HAT*) within populations across the sampling interval using Wilcoxon rank sum test.

| Population Comparison | Test Statistic (W) | P-value |
|---|---|---|
| **ONF Historic - ONF Contemporary** | **440** | **1.47E-03** |
| **ABS Historic - ABS Contemporary** | **80** | **2.74E-08** |
| **JDSP Historic - JDSP Contemporary** | **50** | **3.08E-08** |
| **PLE Historic - PLE Contemporary** | **65** | **7.84E-07** |
| **SPSP Historic - SPSP Contemporary** | **6** | **2.29E-06** |

**Table S11.** Test for significant differences in mean relatedness (*PI_HAT*) between historic populations groups using Wilcoxon rank sum two-tailed test, adjusted for multiple testing.

| Population Comparison | Test Statistic (W) | p-value | Bonferroni-adjusted p-value |
|---|---|---|---|
| **ABS Historic - ONFvONF** | **459** | **3.00E-03** | **3.30E-02** |
| **PLEvPLE_SPSPvSPSP** | **381** | **5.43E-05** | **5.43E-04** |
| **ABSvABS_PLEvPLE** | **93** | **1.05E-06** | **1.05E-05** |
| **ONFvONF_PLEvPLE** | **34** | **2.93E-07** | **2.93E-06** |
| ABSvABS_JDSPvJDSP | 182 | 7.22E-01 | 1.00E+00 |
| ABSvABS_SPSPvSPSP | 292 | 3.78E-01 | 1.00E+00 |
| JDSPvJDSP_ONFvONF | 228 | 1.70E-02 | 1.70E-01 |
| JDSPvJDSP_PLEvPLE | 90 | 1.20E-02 | 1.20E-01 |
| JDSPvJDSP_SPSPvSPSP | 147 | 4.42E-01 | 1.00E+00 |
| ONFvONF_SPSPvSPSP | 123 | 4.30E-02 | 4.28E-01 |

**Table S12.** One-tailed Wilcoxon rank sum test to statistically test the hypothesis that larger populations (ONF and ABS) have lower relatedness (*PI_HAT*) than small populations (PLE, JDSP, SPSP) in contemporary samples. Tests are adjusted for multiple testing.

| Population Comparison | Test Statistic (W) | p-value | Bonferroi-adjusted p-value |
|---|---|---|---|
| **ONF Contemporary < JDSP Contemporary** | **17** | **3.26E-12** | **9.78E-12** |
| **ONF Contemporary < SPSP Contemporary** | **0** | **2.36E-06** | **7.09E-06** |
| **ONF Contemporary < PLE Contemporary** | **0** | **1.72E-09** | **5.16E-09** |
| **ABS Contemporary < JDSP Contemporary** | **126** | **4.88E-11** | **1.46E-10** |
| **ABS Contemporary < SPSP Contemporary** | **14** | **1.05E-06** | **3.14E-06** |
| **ABS Contemporary < PLE Contemporary** | **33** | **6.03E-10** | **1.81E-09** |

**Table S13.** Genes underlying ROH hotspots and cold spots.

| Chromosome | Start | End | Mean ROH Density (Mb) | Length (Mb) | Type | Genes |
|---|---|---|---|---|---|---|
| 1 | 13000001 | 13200000 | 11.89 | 0.2 | hotspot | |
| 1 | 13400001 | 13600000 | 11.96 | 0.2 | hotspot | NCAM2 |
| 1 | 14000001 | 14200000 | 11.53 | 0.2 | hotspot | |
| 1 | 14400001 | 15400000 | 17.37 | 1 | hotspot | App, Atp5pf, GABPA, JAM2, MRPL39 |
| 1 | 16600001 | 18400000 | 13.78 | 1.8 | hotspot | Abhd10, anapc15, Arap1, ARRB1, BACH1, C21orf62, C3orf85, CCT8, CD200, CD200, CD96, CLPB, EMSY, Folr1, Il18bp, Inppl1, LAMTOR1, LRRC32, LTN1, Map3k7cl, NECTIN3, NUMA1, PAXBP1, PDE2A, PHLDB2, Phox2a, Plcxd2, Rnf121, Rwdd2b, STARD10, TAGLN3, THAP12, TMPRSS7, TOMT, TTMP, USP16, Wdr73, XNDC1N |
| 1 | 37800001 | 38000000 | 13.04 | 0.2 | hotspot | g806 |
| 1A | 7400001 | 7600000 | 14.32 | 0.2 | hotspot | KIAA0930, NUP50 |
| 2 | 1600001 | 1800000 | 13.24 | 0.2 | hotspot | LY6E, LY6E, RHPN1, TOP1MT |
| 2 | 70800001 | 71000000 | 11.48 | 0.2 | hotspot | |
| 2 | 130200001 | 130400000 | 11.85 | 0.2 | hotspot | dll1, Dlx5, SDHAF3 |
| 2 | 131000001 | 131800000 | 12.50 | 0.8 | hotspot | acdh-6, Bet1, CASD1, Gng11, PPP1R9A, Sgce, TFPI2 |
| 3 | 97200001 | 97400000 | 11.85 | 0.2 | hotspot | ADI1, eipr1, Rnaseh1, Rps7, Trappc12 |
| 4 | 31000001 | 31200000 | 11.67 | 0.2 | hotspot | |
| 4 | 63400001 | 63600000 | 12.68 | 0.2 | hotspot | |
| 4 | 66000001 | 66200000 | 11.49 | 0.2 | hotspot | |
| 4A | 8800001 | 9000000 | 11.63 | 0.2 | hotspot | Il1rapl2, MAP4K4, RADX |
| 5 | 14800001 | 15200000 | 12.61 | 0.4 | hotspot | DDX24, g13917, g13919, GSC, PPP4R4, Serpina1, SERPINA1, SERPINA10, SERPINA3-4 |
| 5 | 16600001 | 17000000 | 11.80 | 0.4 | hotspot | calm2-b, NRDE2, Psmc1, RPS6KA5, TTC7B |
| 5 | 18600001 | 19000000 | 12.22 | 0.4 | hotspot | FLRT2 |
| 5 | 57600001 | 58000000 | 13.89 | 0.4 | hotspot | ARL14EP, FSHB, KCNA4, MPPED2 |
| 5 | 60600001 | 61000000 | 12.66 | 0.4 | hotspot | Dbx1, NAV2, NELL1, PRMT3 |
| 7 | 24000001 | 24200000 | 14.33 | 0.2 | hotspot | |
| 8 | 27200001 | 27800000 | 13.33 | 0.6 | hotspot | C1orf21, COLGALT2, EDEM3, HMCN1, Ivns1abp, NIBAN1, Rnf2, SWT1, TRMT1L, Tsen15 |
| 8 | 28000001 | 28200000 | 11.57 | 0.2 | hotspot | PLA2G4A, PTGS2 |
| 8 | 31600001 | 31800000 | 13.04 | 0.2 | hotspot | Ier5, KIAA1614, STX6, XPR1 |
| 9 | 15400001 | 15600000 | 14.07 | 0.2 | hotspot | DNER, PID1, Trip12 |
| 15 | 2400001 | 2600000 | 13.02 | 0.2 | hotspot | |
| 1A | 23400001 | 23600000 | 2.66 | 0.2 | coldspot | Cbx6, CBX7, Cby1, Ddx17, DMC1, DNAL4, ENTR1, FAM20C, Gtpbp1, JOSD1, KDELR3, Nptxr, Sun2, TOMM22 |
| 1A | 68200001 | 68600000 | 2.54 | 0.4 | coldspot | CDC123, CELF2, dhtkd1, Echdc3, NUDT5, Proser2, Sec61a2, Upf2, USP6NL |
| 1A | 69600001 | 69800000 | 2.59 | 0.2 | coldspot | |
| 1A | 70200001 | 70400000 | 2.68 | 0.2 | coldspot | ATP5F1C, ITIH2, Itih5, Kin, SFMBT2, STIMATE, TAF3 |
| 2 | 600001 | 1000000 | 2.41 | 0.4 | coldspot | did not map |
| 2 | 8400001 | 8600000 | 2.37 | 0.2 | coldspot | dnaaf11, PHF20L1, TMEM71 |
| 3 | 114000001 | 115000000 | 2.37 | 1 | coldspot | did not map |
| 4 | 1800001 | 2000000 | 2.46 | 0.2 | coldspot | did not map |
| 6 | 33800001 | 34200000 | 2.43 | 0.4 | coldspot | ASAH2, Atad1, LIPA, LIPK, LIPM, MINPP1, PAPSS2, PTEN, RNLS, SGMS1 |
| 7 | 34600001 | 34800000 | 2.62 | 0.2 | coldspot | DUSP19, FRZB, NCKAP1, NUP35 |
| 9 | 23600001 | 23800000 | 2.54 | 0.2 | coldspot | AQP12B, CEP19, g15966, Pak2, PIGX, RYK, Slco2a1, Srprb |
| 11 | 19400001 | 19600000 | 2.42 | 0.2 | coldspot | CA7, CIAO2B, g2137, g2138, PDCD2L, PDP2, RRAD, UBA2, wtip |
| 12 | 1 | 200000 | 2.61 | 0.2 | coldspot | FGD5 |
| 13 | 2000001 | 2200000 | 2.41 | 0.2 | coldspot | NEURL1B, Sh3pxd2b, STK10, UBTD2 |
| 14 | 1 | 200000 | 2.28 | 0.2 | coldspot | GLYR1, MAPK8IP3, MRPS34, ROGDI |
| 15 | 8600001 | 8800000 | 2.50 | 0.2 | coldspot | EP400, NOC4L, Pus1, Susd2, ULK1 |
| 15 | 9000001 | 9200000 | 2.70 | 0.2 | coldspot | MN1, PITPNB, TTC28 |
| 15 | 13200001 | 13400000 | 2.00 | 0.2 | coldspot | SLC15A4, TMEM132C |
| 18 | 5400001 | 5600000 | 2.29 | 0.2 | coldspot | |
| 18 | 7600001 | 8000000 | 2.53 | 0.4 | coldspot | AANAT, Cygb, Exoc7, foxj1, Galr2, JMJD6, Mettl23, MXRA7, PRPSAP1, rhbdf2, RNF157, Sphk1, SRSF2, ST6GALNAC1, ST6GALNAC2, UBALD2, Ube2o |
| 19 | 2800001 | 3000000 | 2.40 | 0.2 | coldspot | Doc2b, g4717, GDPD1, PPM1E, PRR11, RAD51C, ska2, SMG8, TEX14, TRIM37 |
| 19 | 5400001 | 5600000 | 2.37 | 0.2 | coldspot | CCL13, CCL3, CCL8, CCT6, Chchd2, MRPS17, NF2, NIPSNAP2, PHKG1, PSPH, SUMF2, VKORC1L1 |
| 21 | 1200001 | 1400000 | 2.51 | 0.2 | coldspot | HES1, KLHL17, Noc2l, Perm1, Plekhn1, Samd11 |
| 21 | 1600001 | 2000000 | 2.29 | 0.4 | coldspot | ACAP3, B3galt6, c1qtnf12, CPTP, Dvl1, INTS11, Lpar6, MXRA8, pusl1, scnn1a, SDF4, TAS1R3, TNFRSF18, TNFRSF4, UBE2J2 |
| 21 | 3000001 | 3200000 | 2.70 | 0.2 | coldspot | |
| 22 | 1600001 | 1800000 | 2.18 | 0.2 | coldspot | BIN3, BMP1, C8orf58, EGR3, EGR3, LGI3, Pdlim2, Phyhip, Piwil2, PPP3CC, SFTPC, SFTPC, Slc39a14, SORBS3 |
| 22 | 3000001 | 3200000 | 2.25 | 0.2 | coldspot | |
| 25 | 1200001 | 2000000 | 1.58 | 0.8 | coldspot | did not map |
| 26 | 3600001 | 4000000 | 2.37 | 0.4 | coldspot | APOBEC2, BAK1, Cd300ld, FOXP4, IP6K3, Itpr3, MDFI, Nfya, Oard1, PGB, PGB, Tfeb, TREM2, UQCC2 |
| 27 | 3600001 | 4400000 | 2.22 | 0.8 | coldspot | did not map |
| 27 | 4600001 | 5200000 | 2.33 | 0.6 | coldspot | ABI3, Asb16, Atp5mc1, Atxn7l3, CALCOCO2, COL1A1, Dlx3, dlx4a, G6pc3, Gip, GNGT2, HOXB1, hoxb13a, HOXB2, HOXB3, HOXB4, hoxb5a, Hoxb6, HOXB7, HOXB8, HOXB9, HROB, IGF2BP1, ITGA3, KAT7, LSM12, Mpp2, NGFR, NXPH3, Pdk2, PHB1, PHOSPHO1, Ppp1r9b, PPY, pyya, retsat, Samd14, SGCA, SNF8, SPOP, TMEM101, Tmub2, Ube2z, Znf652 |

**Table S14.** Key GO results: filtered "GO enrichment" to require at least 3 genes mapped to the term (Significant >= 3) and that the genes in the term come from at least two different regions (reg.ct >= 2) and the top "biological process" (BP) terms, one for "hi", one for "low".

| GO type | GO ID | Term | Annotated | Significant | Expected | Classic Fisher | topgo Fisher | Genes | Region count | Type |
|---|---|---|---|---|---|---|---|---|---|---|
| BP | GO:0010466 | negative regulation of peptidase activity | 34 | 6 | 0.25 | 0.00000014 | 0.01368 | g13917, SERPINA1, g13919, Serpina1, SERPINA3-4, TFPI2 | 2 | hotspots |
| MF | GO:0004867 | serine-type endopeptidase inhibitor activity | 57 | 8 | 0.43 | 9.4E-09 | 9.4E-09 | SERPINA10, g13917, SERPINA1, g13919, Serpina1, SERPINA3-4,App, TFPI2 | 3 | hotspots |
| MF | GO:0061630 | ubiquitin protein ligase activity | 159 | 4 | 1.2 | 0.03235 | 0.03235 | LTN1, Rnf121, Rnf2, Trip12 | 3 | hotspots |
| CC | GO:0009986 | cell surface | 166 | 5 | 1.15 | 0.00584 | 0.0015 | CD200, CD200, Folr1, App, JAM2 | 2 | hotspots |
| CC | GO:0031225 | anchored component of membrane | 21 | 3 | 0.15 | 0.00039 | 0.002 | Folr1, LY6E, LY6E | 2 | hotspots |
| CC | GO:0005615 | extracellular space | 607 | 11 | 4.22 | 0.00289 | 0.0029 | LRRC32, NELL1, FSHB, FLRT2, SERPINA10, g13917, SERPINA1, g13919, Serpina1, SERPINA3-4, TFPI2 | 6 | hotspots |
| CC | GO:0043025 | neuronal cell body | 59 | 3 | 0.41 | 0.00791 | 0.0079 | CD200, CD200, HMCN1 | 2 | hotspots |
| CC | GO:0030424 | axon | 116 | 4 | 0.81 | 0.00853 | 0.0085 | CD200, CD200, KCNA4, HMCN1 | 3 | hotspots |
| BP | GO:0009952 | anterior/posterior pattern specification | 36 | 5 | 0.51 | 0.00014 | 0.00014 | HES1, HOXB9, Hoxb6, hoxb5a, HOXB4 | 2 | coldspots |
| MF | GO:0016788 | hydrolase activity, acting on ester bonds | 514 | 14 | 7.28 | 0.01436 | 0.0211 | MINPP1, PTEN, LIPK, LIPM, LIPA, PDP2, PSPH, RAD51C, PPM1E, GDPD1, INTS11, PPP3CC, G6pc3, PHOSPHO1 | 7 | coldspots |
| MF | GO:0005179 | hormone activity | 76 | 4 | 1.08 | 0.02263 | 0.0226 | c1qtnf12, pyya, PPY, Gip | 2 | coldspots |
| MF | GO:0005035 | death receptor activity | 8 | 3 | 0.11 | 0.00015 | 0.0278 | TNFRSF4, TNFRSF18, NGFR | 2 | coldspots |
| MF | GO:0000981 | DNA-binding transcription factor activity, RNA polymerase II-specific | 661 | 16 | 9.36 | 0.02509 | 0.0386 | foxj1, HES1, Nfya, FOXP4, Dlx3, dlx4a, hoxb13a, HOXB9, HOXB8, HOXB7, Hoxb6, hoxb5a, HOXB4, HOXB3, HOXB2, HOXB1 | 4 | coldspots |
| CC | GO:0005576 | extracellular region | 863 | 16 | 12.08 | 0.14985 | 0.00506 | ASAH2, RNLS, FRZB, g15966, Susd2, CCL13, CCL8, CCL3, TEX14, c1qtnf12, SFTPC, SFTPC, pyya, PPY, COL1A1, Gip | 9 | coldspots |
| CC | GO:0009986 | cell surface | 166 | 5 | 2.32 | 0.08342 | 0.01829 | MXRA8, TNFRSF18, TREM2, Cd300ld, NGFR | 3 | coldspots |
| CC | GO:0005741 | mitochondrial outer membrane | 44 | 3 | 0.62 | 0.0234 | 0.0234 | Atad1, TOMM22, BAK1 | 3 | coldspots |
